# A cross-sectional examination of immune adaptations during pregnancy in the ECHO Cohort

**DOI:** 10.64898/2026.02.04.703821

**Authors:** Sarah M. Banker, Rosa Shapiro-Thompson, Sarah Sinsel, Akhgar Ghassabian, Christian Douglas, Morgan E. Nelson, Lisa A. Peterson, Bharat Thyagarajan, Santiago Morales, Christine W. Hockett, Amy J. Elliott, Stephanie N. Giamberardino, Lauren C. Shuffrey, the ECHO Cohort Consortium

## Abstract

**Background:** Pregnancy requires finely tuned immune changes that support implantation, placental development, maternal–fetal tolerance, and preparation for labor, yet the normative trajectories of circulating inflammatory proteins across gestation remain poorly defined. This cross-sectional study investigates how circulating inflammatory proteins vary with gestational age in pregnancy and examines the impacts of fundamental biological characteristics, such as gravidity and fetal sex.

**Methods:** Data were drawn from 1154 pregnant individuals from six study sites of the National Institutes of Health Environmental influences on Child Health Outcomes (ECHO) Cohort. We used Olink high-throughput proteomic profiling to map cross-sectional associations between protein expression levels and gestational age at blood draw using linear, spline-based, and generalized additive modeling approaches.

**Results:** Generalized additive models provided the best fit, revealing that immune changes across pregnancy were predominantly nonlinear. Sixty-one proteins showed significant associations with gestational age, with many exhibiting shared inflection points that aligned with major physiological transitions. A small subset of proteins also showed evidence of modification by fetal and maternal characteristics. CD244 displayed different gestational patterns by fetal sex, while CST5 and SIRT2 showed varied gestational associations by maternal gravidity.

**Conclusion:** The findings highlight pregnancy as a sequence of coordinated immune transitions rather than a simple linear shift and provide one of the most detailed characterizations to date of circulating inflammatory protein dynamics across human gestation. Establishing these normative trajectories offers a crucial reference for detecting early deviations that may signal risk for pregnancy complications and for identifying biomarkers in maternal and fetal health research.

## INTRODUCTION

Over the course of pregnancy, the maternal immune system undergoes significant changes to support the adapting needs of the developing fetus while safeguarding maternal health. A comprehensive understanding of the normative trajectory of these immunological adaptations is critical, as disruptions to this delicate balance have been implicated in pregnancy complications, including preeclampsia (1,2), gestational diabetes (3,4), pregnancy loss (5,6), and preterm birth (7–9).

Understanding how immune functioning aligns with the physiological stages of pregnancy offers critical context for interpreting changes in the expression of specific proteins. Pregnancy involves both innate immunity, which provides rapid responses to changes in the maternal environment, and adaptive immunity, which develops more slowly and confers long-lasting immune regulation (10–12). Innate immunity is mediated by cells such as uterine natural killer (NK) cells and macrophages, while adaptive immunity is driven by regulatory T cells (Tregs) and T helper (Th) cells (10–12). Both innate and adaptive immune mechanisms rely on a diverse array of signaling molecules, including chemokines, cytokines, growth factors, and immunomodulatory receptors, that orchestrate these processes across pregnancy. Although the precise roles of many specific immune-related proteins remain unclear beyond a well-studied subset (e.g., IL-6, IL-8, TNF-α, and IFN-γ), there is a growing understanding of how broader groups of immune mediators support pregnancy.

During the peri-implantation period and early first trimester, the maternal uterine environment adopts a controlled pro-inflammatory profile to support embryo implantation and early placental development. Innate immune cells accumulate in the decidua and initiate blastocyst attachment, trophoblast invasion, and early vascular remodeling (12–15). Chemokines and classical inflammatory cytokines, including IL-6 and IL-8, recruit leukocytes and promote the localized extracellular matrix breakdown needed for invasion (16,17). A brief, limited Th1-type signal also contributes during this window, supporting trophoblast invasion before being rapidly downregulated (18). Together, these coordinated innate and adaptive signals establish the conditions necessary for uteroplacental blood flow (19). Shortly after implantation, Th1 immunity is replaced by Th2/Treg-mediated anti-inflammatory responses, which promote maternal–fetal tolerance and create a tolerogenic environment that supports placental and fetal growth (11,12,20). Chemokines play an important role in this adaptive immunity by shaping T cell responses, while immunoregulatory receptors such as PD-L1 contribute to maternal–fetal tolerance and maintain an environment that protects the fetus from rejection (18,21). This anti-inflammatory state extends into the second and third trimesters, accompanied by finely tuned immune modulation and controlled changes in specific mediators that safeguard both maternal and fetal health (22). Near term, the immune environment transitions back to a pro-inflammatory profile, driven primarily by innate immune mechanisms that stimulate uterine contractility and cervical ripening (22,23).

Pregnancy is often described in terms of these broad immune shifts, such as early pro-inflammatory activation followed by mid-pregnancy immune tolerance. In line with this framework, most empirical studies evaluate gestational differences using linear or trimester-based comparisons. This approach implicitly assumes smooth, monotonic change across pregnancy and may obscure biologically meaningful inflection points that coincide with well-established physiological transitions, such as the onset of placental perfusion near the end of the first trimester or the pre-labor inflammatory surge. Only a handful of studies have examined whether circulating immune markers follow non-linear or stage-specific trajectories (24–28), and comprehensive mapping of these patterns has remained limited by small cohorts, targeted panels, and sparse sampling across gestation. A more flexible, data-driven characterization of immune change is therefore needed to identify when, rather than simply whether, specific immune mediators are most active.

In this study, we aimed to comprehensively describe how circulating inflammatory proteins vary with gestational age (GA) in pregnancy. To do so, we leveraged the breadth of the National Institutes of Health (NIH) Environmental influences on Child Health Outcomes (ECHO) Cohort, which enrolled pregnancies across diverse U.S. populations. Importantly, we did not restrict our sample to exclusively healthy pregnancies; instead, we sought to characterize patterns of immune protein expression across gestation in a way that reflects the range of maternal backgrounds and conditions encountered in the general population. We also sought to examine the impacts of fundamental biological characteristics, such as gravidity, as prior pregnancies are thought to “train” the maternal immune system (29–31), and fetal sex, as pregnancies carrying male fetuses are thought to display more pro-inflammatory maternal immune responses (32,33). By taking this approach, we aim to establish a baseline characterization of gestational immune trajectories that can serve as a reference point for future investigations into how specific maternal health factors, complications, or exposures may alter these processes.

## MATERIALS AND METHODS

### Study Design

Data for this analysis were collected from participants recruited at six study sites of the ECHO Cohort, which consists of longitudinal birth cohort studies across the U.S. and Puerto Rico. The goal of the ECHO Program is to investigate how early-life environmental exposures affect child health and development across five primary domains: pre-, peri-, and postnatal outcomes; upper and lower airway conditions; obesity; neurodevelopment; and positive health (34–37).

Participants were selected for this assay if they had pregnancy plasma samples banked in the ECHO biorepository, had no limitations on use of biospecimens noted, were part of a site that had at least 50 participants with eligible samples, and were enrolled in ECHO sites that collected information on pregnancy health conditions and administered mental health assessments during the perinatal period (N=1606). All participants consented to participate in their local ECHO site and share their information with the ECHO consortia. Both a central and site-specific Institutional Review Board monitored human subject activities at each site and the centralized ECHO Data Analysis Center. Written informed consent and the parent’s/guardian’s permission were obtained, along with child assent as appropriate, for ECHO-wide Cohort Data Collection Protocol participation. Reporting followed the Strengthening the Reporting of Observational Studies in Epidemiology (STROBE) guidelines for reporting observational studies.

### Maternal Venous Blood Collection

Maternal venous blood was collected by trained phlebotomy staff into ethylenediaminetetraacetic acid-treated vacutainer tubes at a single study visit between 5 and 41 weeks’ gestation. Immediately after collection, tubes were gently inverted 8–10 times to ensure anticoagulation and transported on wet ice to the processing laboratory within 2 hours. Blood samples were centrifuged at 1500 × g for 10 minutes at 4°C to separate plasma. The plasma supernatant was carefully aspirated without disturbing the buffy coat or red blood cell layer and aliquoted into pre-labeled cryovials (120 μL per aliquot) using sterile technique. Aliquots were immediately flash-frozen on dry ice and subsequently transferred to -80°C long-term storage freezers, then shipped to the ECHO central biorepository, Fisher Bioservices. Once aliquots were selected for analysis, 40 μL were realiquoted into 96-well plates and subsequently shipped on dry ice to the NIH-funded Minnesota Human Health Exposure Analysis Resource (HHEAR) Laboratory for biomarker assays.

### Proteomic Profiling and Data Cleaning

Olink Target 96 Inflammation panel data, which quantified 92 immune-related proteins in plasma using proximity extension assay technology, were generated for a total of 1606 samples. Each protein is expressed using a Normalized Protein eXpression (NPX) unit, which is a log2-transformed value representing the relative abundance of each protein. Panel quality control and intensity normalization procedures were conducted by the data-generating lab according to manufacturer guidelines. Two samples failed during data generation and were immediately excluded from any analysis. To identify samples that were potential outliers compared to the majority of samples, we followed procedures mirroring those applied by the UK Biobank (38) and excluded samples whose median NPX, interquartile range, and principal component 1 or principal component 2 (the two principal components capturing the largest amount of variance in the dataset) were >5 standard deviations from the dataset mean, resulting in N=4 samples being flagged for exclusion from association testing. Concurrently, using the N=1604 samples as input, the proportion of samples with an NPX value ≤ the limit of detection (LOD) (set as the maximum LOD across all assay plates) was calculated for each protein, and proteins with a proportion of samples ≤ LOD < 0.25 were retained for analysis (N=69). Finally, among N=57 technical replicate pairs (N=114 total samples), one was chosen at random to remain in the analysis and the other was excluded from downstream association testing (see Supplementary Table 1 for the coefficients of variation). Samples flagged with a warning for having a control probe >0.3 NPX from a plate’s median remained in the analysis (N=122 of 1543 samples). In total, 1543 unique samples (1502 individuals, 1543 pregnancies) and 69 proteins (Supplementary Table 2) passed proteomic data cleaning and were eligible for downstream analysis.

### Current Analysis

Participants in the current analysis included pregnant participants with valid Olink proteomic data, confirmed singleton pregnancies, and available data on GA at blood draw. Those with GA <4 weeks or >41 weeks were excluded for plausibility, yielding N=1354 participants. To ensure independence of observations, participants who had proteomic profiling data available from more than one pregnancy (N=30 participants, N=62 pregnancies) were restricted to the first available pregnancy; data from subsequent pregnancies from the same individual were excluded from analysis (see Supplementary Figure 1 for an analysis of protein correlations across multiple pregnancies). We additionally excluded participants whose pregnancy was missing information on gravidity (N=159) and child sex (N=9), as these are crucial covariates known to impact the prenatal inflammatory environment (29–33). After exclusions, a total of 1154 participants were included in the current analysis (see Supplementary Figure 2 for a flow chart outlining sample exclusions).

GA at blood draw was calculated using the difference between the expected due date and specimen collection date. Specifically, we computed the number of days between the blood draw and the expected due date, converted this value to weeks, and subtracted the result from 40 weeks (full-term). Gravidity was defined as the number of times a participant had been pregnant, including the current pregnancy, regardless of the viability outcomes. Fetal sex was obtained from information on the child’s sex at birth. Information on expected due date, gravidity, and child’s sex at birth was extracted from medical records or self-reported by participants.

### Statistical Analysis

#### Inflammatory associations with gestational age

To identify markers associated continuously with GA, we modeled each protein separately, controlling for maternal age at blood draw, gravidity, and infant sex and accounting for heterogeneity across sites by including a random intercept for ECHO site of enrollment. Gravidity was categorized as first pregnancy, second pregnancy, or third or later pregnancy. To compare linear and non-linear fits for the relationships between GA and protein levels, we initially examined three model families: linear mixed-effects models, spline-based mixed-effects models, and generalized additive models (GAMs). Non-linear models were included given prior evidence that immune adaptation across pregnancy may follow non-linear trajectories, with accelerated or plateauing changes in later gestation. The linear models included GA as a continuous fixed effect. The spline models used natural cubic splines with three degrees of freedom to allow moderate flexibility in the GA–protein association. GAMs were fit using penalized regression splines with a smooth term for GA (estimated by restricted maximum likelihood) and a random-effect smooth for ECHO site of enrollment. For each GAM, we used the Wald F-test from the mgcv package (39) in R to assess whether the smooth for GA differed significantly from a flat (no-effect) line; if mgcv-reported p-values were truncated to zero due to precision limits within the function, they were recomputed manually from the same F-statistic and degrees of freedom. For each model, we extracted the estimated degrees of freedom (EDF) (where applicable), test statistics for the GA term (t-statistic for linear models and F-statistic for spline models and GAMs), and model fit indices including the Akaike information criterion (AIC), marginal R², and conditional R². To evaluate which functional form best described protein–GA relationships, we compared AIC values across model families for each protein and identified the model with the lowest AIC as the best fit. All p-values for GA effects were corrected for multiple testing within each model family using the Benjamini–Hochberg false discovery rate (FDR) procedure at 5%. For easy comparison of our findings with prior work, we also evaluated trimester-level differences in protein expression levels (see Supplementary Figure 3 and Supplementary Table 3).

#### Effect modification by fetal sex, gravidity, or maternal age

To test whether GA associations differed by key effect modifiers, we compared nested GAMs using maximum likelihood-based joint tests. For categorical modifiers (fetal sex and gravidity), a baseline model including a single smooth for GA (shared across all participants) was compared to a full model that additionally allowed modifier-specific smooths for GA (e.g., separate curves by sex or gravidity group). For the continuous modifiers (maternal age at blood draw), the baseline model included main-effect smooths for GA and maternal age, and the full model additionally included a tensor-product interaction smooth to capture continuous modification of GA trajectories by maternal age. All models adjusted for the remaining covariates (maternal age, gravidity, and fetal sex, as appropriate) and included a random-effect smooth for ECHO site.

Smooth terms were represented using thin-plate regression splines with a modest basis dimension (k=5) to balance flexibility and interpretability, and basis adequacy was confirmed via the mgcv diagnostic k-index (40). Model comparison was performed using likelihood-ratio tests, providing a single F-statistic and p-value reflecting the joint evidence that GA–protein trajectories differed across modifier levels (for the categorical modifier) or continuously by maternal age (for the continuous modifier).

For categorical modifiers (fetal sex and gravidity), significant interactions indicate proteins for which the model with separate GA smooths for each group provided a significantly better fit than a model with a single, shared smooth across all participants. For the continuous modifier (maternal age at blood draw), significant interactions indicate proteins for which adding a tensor-product interaction smooth improved model fit relative to a model with only main-effect smooths for GA and maternal age, reflecting continuous modification of GA trajectories by maternal age. P-values were adjusted across proteins for each modifier using the Benjamini–Hochberg FDR procedure. Because interaction tests generally have lower statistical power than main-effect tests, we applied a more lenient FDR threshold (10%).

In a small subset of proteins, the full and baseline models converged to nearly identical fits, yielding zero or negative model deviance differences (i.e., no additional variance explained by the modifier-specific smooths). For these markers, the interaction was considered non-significant, and results were reported as NA to reflect the absence of a valid comparison. For categorical modifiers that showed significant interactions, we conducted additional sensitivity analyses to confirm that apparent group differences were not driven by uneven sampling across GA. Specifically, we compared the overall GA distributions between groups using global density plots and chi-square tests on GA-by-trimester contingency tables.

To evaluate whether overall protein expression levels were related to key maternal and fetal characteristics independent of their associations with GA, we additionally fit linear mixed-effects models testing main effects of fetal sex, gravidity, and maternal age on expression level for each protein. All models adjusted for the other covariates (maternal age, gravidity, and fetal sex, as appropriate) and included a random intercept for ECHO site. The results are provided in the Supplementary Materials (Supplementary Figure 4; Supplementary Tables 4–6).

## RESULTS

Sample characteristics are presented in Table 1. Maternal age at blood draw was an average of 30 years and ranged from 15 to 44 years. Participants were recruited from six different ECHO sites that spanned recruitment across eight U.S. states. GA at blood draw spanned nearly the full course of pregnancy, ranging from 5 to 40 weeks, though the majority of samples were collected in the third trimester. This distribution highlights the strength of the study in capturing late pregnancy at scale, while still including a sizable number of early- and mid-pregnancy samples. Fetal sex was approximately evenly distributed. Gravidity was also well represented, with roughly 22% of mothers in their first pregnancy, 34% in their second, and 44% in their third or later pregnancy.

**Table 1.** Characteristics of the participants (N=1154)

| Characteristic | Mean (sd, range) or Count (%) |
| --- | --- |
| Maternal age at blood draw (years) | 30.71 (4.90, 15.23–44.5) |
| Maternal race | American Indian or Alaska Native: <25 (<2%)<br>Asian: 35 (3.03%)<br>Black: 189 (16.38%)<br>Native Hawaiian or other Pacific Islander: <15 (<2%)<br>White: 790 (68.46%)<br>Multiple races: 35 (3.03%)<br>Other race: 37 (3.21%) |
| Maternal ethnicity | Hispanic: 140 (12.13%)<br>Not Hispanic: 1010 (87.52%)<br>Inconsistent or no information: 4 (0.35%) |
| Gestational age at blood draw (weeks) | 30.65 (8.37, 5.71–40.86) |
| Trimester of blood draw | 1<sup>st</sup> trimester: 103 (8.93%)<br>2<sup>nd</sup> trimester: 169 (14.65%)<br>3<sup>rd</sup> trimester: 882 (76.43%) |
| Fetal sex | Male: 564 (48.87%)<br>Female: 590 (51.13%) |
| Gravidity | 1<sup>st</sup> pregnancy: 254 (22.01%)<br>2<sup>nd</sup> pregnancy: 390 (33.80%)<br>3<sup>rd</sup>+ pregnancy: 510 (44.19%) |
| Site (recruitment location; years) | Safe Passage Study (Sioux Falls and Rapid City, South Dakota; 2019-2023): 401 (34.75%)<br>Children's Respiratory and Environmental Workgroup (CREW 3; Detroit, Michigan; Madison, Wisconsin, Nashville, Tennessee; St. Louis, Missouri; 2020–2023): 243 (21.06%)<br>Pittsburgh Girls Study (Pittsburgh, Pennsylvania; 2019–2022): 85 (7.37%)<br>Michigan Archive for Research on Child Health (MARCH; East Lansing, Michigan; 2019–2023): 176 (15.25%)<br>The NYU Children's Health and Environment Study (NYU CHES; New York, NY; 2019-2022): 105 (9.10%)<br>PRogramming of Intergenerational Stress Mechanisms (PRISM; Boston, MA; New York, NY; 2019-2022): 144 (12.48%) |

### Inflammatory Associations with Gestational Age

Across proteins, GAMs most frequently provided the best model fit, yielding the lowest AIC for 67 of 69 proteins (97.1%). Only two proteins were better fit by spline models, and none were best fit by linear models. Mean and median AIC values were also lowest for GAMs (mean=2106, median=1992) compared to linear models (mean=2138, median=2006) and spline-based mixed-effects models (mean=2122, median 2006). On this basis, GAMs were selected as the primary analytic approach (see Supplementary Tables 7–9 for the full modeling results). We identified 61 inflammatory proteins out of 69 that showed significant associations with GA after correcting for multiple comparisons with an FDR of 5% (Figure 1).

**Figure 1.**
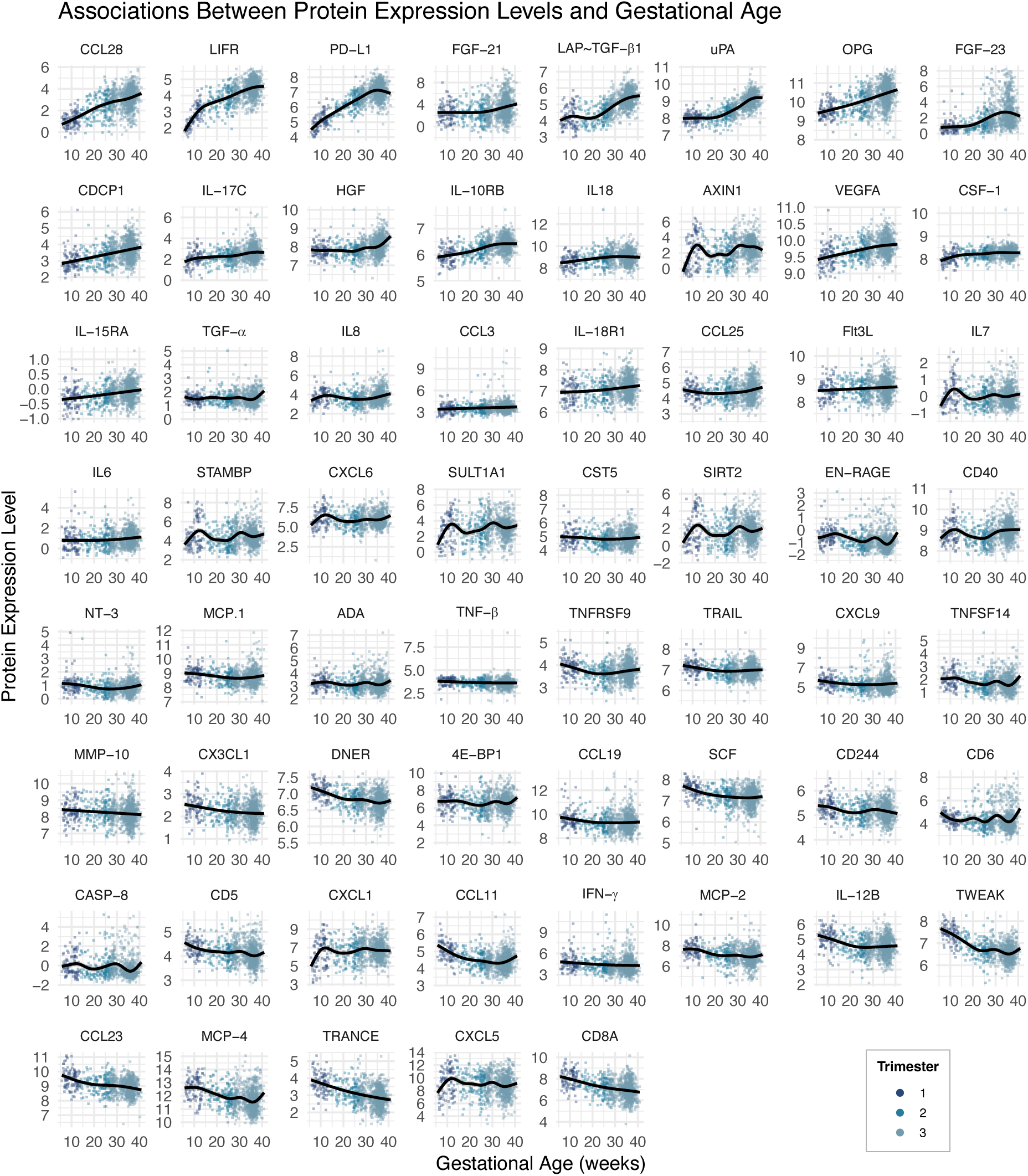
Associations between protein expression levels and gestational age, sorted by beta estimate. Univariate models detected 61 inflammatory proteins that showed significant associations with gestational age at 5% false discovery rate. Generalized additive models controlled for maternal age at blood draw, gravidity, and infant sex and included site as a random effect. Cases are labeled by trimester of blood draw collection.

Notably, a subset of proteins displayed a shared and distinct non-linear pattern with pronounced inflections around approximately 12 and 30 weeks of gestation: SIRT2, AXIN1, STAMBP, EN-RAGE, IL-7, CASP-8, and CXCL5.

### Modification Effects of Fetal Sex, Gravidity, or Maternal Age

One protein, CD244, exhibited a significant difference in GA trajectories by fetal sex (F(4.05, 1138.69)=5.13, p-FDR=0.03; Figure 2). Follow-up smooth-term analyses indicated that this effect was driven by a nonlinear pattern among participants pregnant with female fetuses, who showed a significant dip in early to mid-gestation followed by stabilization later in pregnancy (EDF=3.05, F=5.59, p<0.001), whereas trajectories among those pregnant with male fetuses were largely flat across gestation (p=0.99). Two proteins, CST5 and SIRT2, exhibited significant differences in GA trajectories by gravidity group (CST5: F(0.30, 1139.82)=26.50, p-FDR=0.05; SIRT2: F(4.81, 1137.89)=4.03, p-FDR=0.05; Figure 3). Follow-up smooth-term analyses indicated that these effects were driven primarily by nonlinear patterns, with early gestational declines followed by stabilization later in pregnancy among multigravida (3+ pregnancies) participants (CST5: EDF=2.86, F=3.72, p=0.02; SIRT2: EDF=3.18, F=4.68, p=0.001), whereas trajectories were largely flat in participants experiencing their first and second pregnancies (p values >0.30). No significant GA × maternal age interactions were detected after correction for multiple comparisons (FDR<0.10), indicating that the overall patterns of immune change across gestation were broadly similar across the maternal age range represented in the sample (see Supplementary Tables 10–12 for full interaction modeling results).

**Figure 2.**
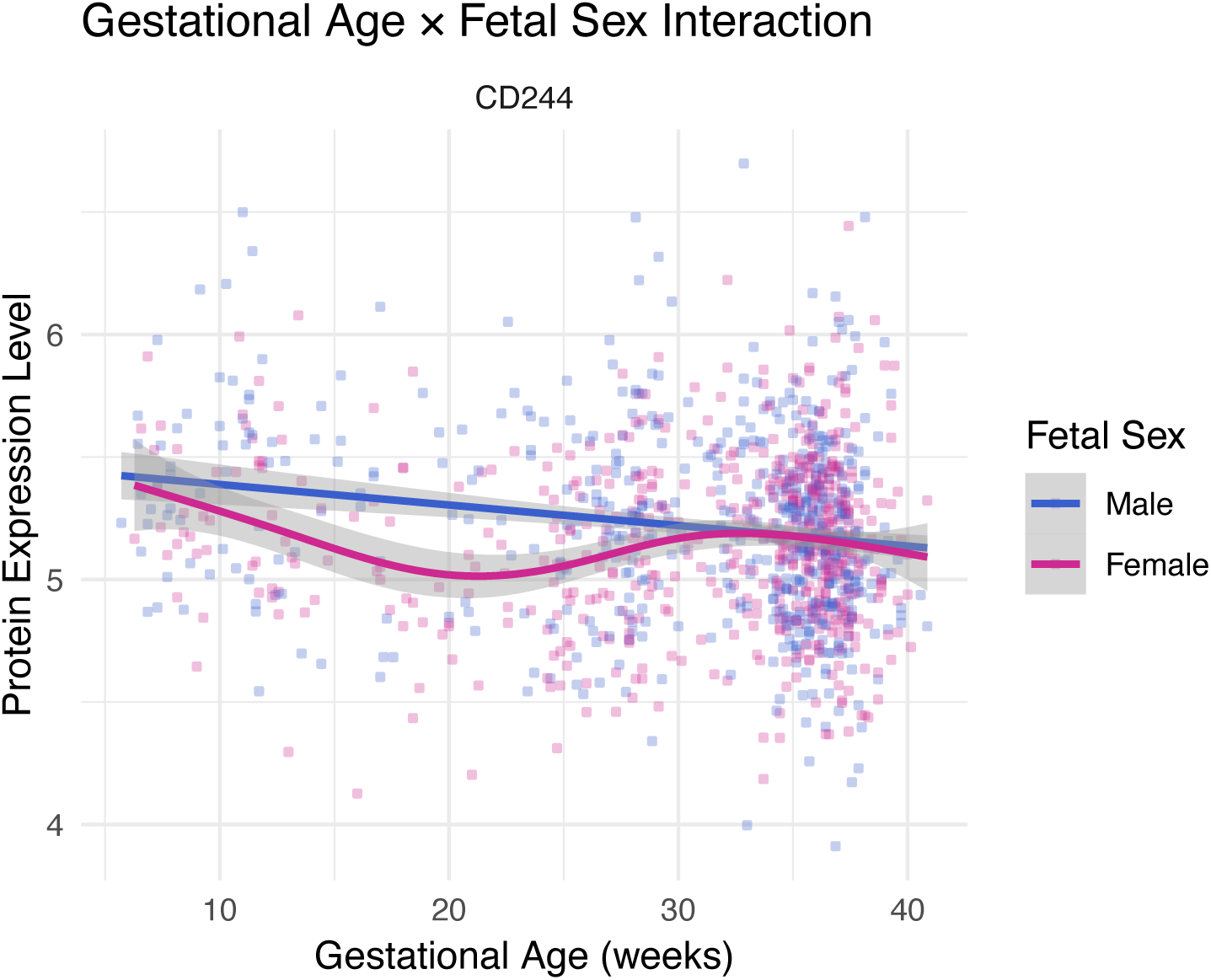
Interaction between gestational age and fetal sex on CD244 expression level. Smoothed curves illustrate unadjusted relationships between gestational age and CD244 protein expression levels (Normalized Protein eXpression, NPX) by fetal sex. Participants pregnant with female fetuses exhibited an early-to-mid-gestational decline followed by stabilization later in pregnancy, whereas trajectories among male fetuses remained relatively flat across gestation. Shaded bands represent 95% confidence intervals from the fitted generalized additive model curves. Full model results are provided in the Supplementary Materials.

**Figure 3.**
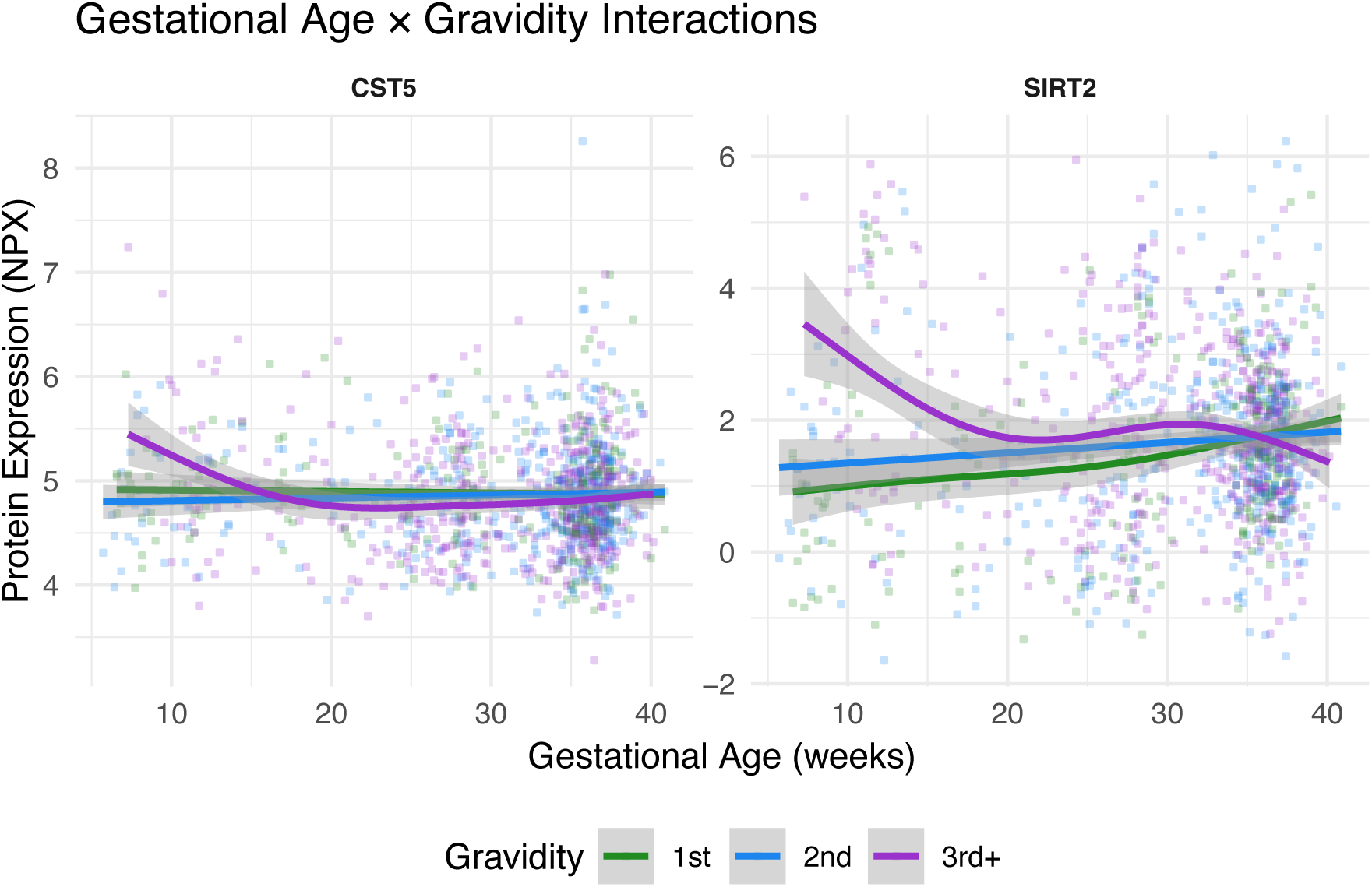
Interactions between gestational age and maternal gravidity on protein expression level. Smoothed curves illustrate unadjusted relationships between gestational age and CST5 and SIRT2 protein expression levels (Normalized Protein eXpression, NPX) within each gravidity group. Both proteins showed early gestational declines followed by stabilization later in pregnancy among multigravida (3+ pregnancies) participants, whereas trajectories were largely flat among primigravida and secundigravida participants. Shaded bands represent 95% confidence intervals from the fitted generalized additive model curves. Full model results are provided in the Supplementary Materials.

To ensure that the observed fetal sex- and gravidity-related effects were not driven by unequal sampling across GA, we compared the GA distributions between groups. For fetal sex, a global Kolmogorov–Smirnov test indicated no overall differences in GA sampling (D<0.05, p>0.30), and a chi-square test of GA-by-trimester frequencies likewise revealed no significant imbalance between male and female fetuses across trimesters (χ²(2)<3.0, p>0.10). These results suggest that the observed divergence in CD244 trajectories by fetal sex reflects genuine biological variation rather than sampling bias. For gravidity, a global Kolmogorov–Smirnov test indicated no overall distributional differences (D=0.08, p=0.29). However, a chi-square test on

GA-by-trimester frequencies revealed modest differences across trimesters (χ2(4)=10.71, p=0.03), with later-trimester samples slightly overrepresented among multigravida (3+ pregnancies) participants. Importantly, when the comparison was restricted to the first trimester, where modeled group differences were most pronounced, no significant differences in sampling frequency were observed (χ2(2)=0.37, p=0.83). Such results indicate that the observed early gestational divergence in trajectories was not attributable to uneven sampling.

## DISCUSSION

In this large multi-site investigation, we found significant gestational changes across a broad network of inflammation-related proteins. GAMs provided the best fit for nearly all proteins examined (97% of 69 proteins), underscoring that immune changes across pregnancy follow mostly non-linear patterns, including phase-like transitions aligned with key physiological milestones. These findings extend prior work that has characterized broad changes by providing a more refined, continuous map of immune signaling from early implantation through term. Importantly, our dataset spans nearly the entire course of human pregnancy, capturing both very early (<6 weeks) and late (>40 weeks) gestational windows that have historically been underrepresented in human studies.

We identified 61 proteins that showed significant associations with GA using flexible non-linear modeling. The observed trajectories align with established prenatal immune frameworks. Proteins involved in early immune activation, including TWEAK, TRANCE, MCP-4, CD8A, and CCL11, were elevated early and declined thereafter, likely reflecting the transient pro-inflammatory signaling necessary for implantation and early placental remodeling (41–44). In contrast, proteins supporting tissue remodeling and vascular maintenance, such as LIF-R, uPA, and OPG, showed progressive increases across gestation, consistent with sustained extracellular matrix remodeling and the preservation of placental and vascular integrity (45–47). Concurrently, mediators of immune tolerance and anti-inflammatory regulation, including PD-L1, LAP–TGFβ1, IL-10RB, and CCL28, also rose over gestation, likely supporting the establishment and maintenance of maternal–fetal immune tolerance (48–52). Within these general trajectories, the models revealed biologically meaningful non-linearities. For instance, PD-L1 exhibited a modest dip toward the end of pregnancy after a steady rise through mid-gestation, potentially reflecting a reduced need for immune suppression as labor approaches. LAP–TGFβ1 and uPA showed delayed increases beginning after approximately 20 weeks, suggesting stage-specific activation of tolerogenic and remodeling pathways. Upon visual inspection, several proteins, including SIRT2, AXIN1, STAMBP, EN-RAGE, IL-7, CASP8, and CXCL5, also displayed pronounced non-linear inflections around approximately 12 and 30 weeks of gestation. These timepoints correspond to major immunological and physiological transitions in pregnancy: namely, the establishment of placental perfusion near the end of the first trimester and the metabolic and immune activation that precedes labor in the third (11,53).

Gestational immune trajectories were further modulated by fetal sex and maternal reproductive history. The cell-surface receptor CD244, expressed on both NK and T cells, exhibited a striking sex-specific trajectory. Pregnancies with female fetuses showed a decrease in CD244 expression during early-to-mid gestation, followed by an increase during late pregnancy, whereas trajectories among male fetuses remained relatively flat. CD244 serves as a co-regulatory receptor that can transmit inhibitory or activating signals depending on cellular context (54,55). In decidual NK cells, it predominantly mediates inhibitory signaling (56). The observed U-shaped expression among pregnancies with female fetuses may reflect transient adjustment of activation thresholds during immune adaptation. This interpretation aligns with recent single-cell studies demonstrating that uterine NK cells undergo continuous ligand- and hormone-driven differentiation across gestation, accompanied by dynamic remodeling of their inhibitory and activating receptor repertoire (57). Such coordinated shifts could underlie the observed sex-specific modulation of CD244 expression. Consistent with this framework, female fetuses have been shown to elicit more precisely tuned maternal immune regulation, with inflammatory pathways adjusting in a more targeted and context-dependent manner across gestation (32,33). In contrast, pregnancies carrying male fetuses display stronger inflammatory responses (32,33). Given that dysregulation of NK cell signaling and cytotoxic receptor expression has been implicated in recurrent pregnancy loss, implantation failure, and preeclampsia (58–61), this sexually dimorphic pattern in CD244 may represent a mechanism through which fetal sex contributes to differential susceptibility to pregnancy complications (62). Longitudinal and cell type-specific studies will be needed to determine whether these circulating patterns reflect localized modulation within the decidua or broader systemic immune tuning.

We also observed that participants with multiple prior pregnancies showed higher early gestational levels of CST5 and SIRT2 expression, followed by stabilization later in pregnancy. CST5 encodes cystatin D, which suppresses enzymes that digest extracellular matrix proteins, such as cathepsins. While cystatin D has not been widely studied in pregnancy, its function as a cysteine-protease inhibitor suggests it could limit the extent of tissue breakdown and remodeling during placental development (63). SIRT2 is a deacetylase that modifies metabolic and inflammatory regulators, helping cells balance energy production and immune activation (64,65). In pregnancy, SIRT2 is expressed in decidual and trophoblast cells, where it may help support placental development by restraining excessive inflammatory and oxidative stress (66,67). The trajectories observed in third and subsequent pregnancies are consistent with the concept of a “trained” uterine immune environment, wherein prior pregnancies condition decidual and vascular tissues to mount a more rapid or contained regulatory response upon subsequent pregnancies (29–31). This priming effect may reduce the need for intense early tissue remodeling signals in multigravida individuals, preventing excessive inflammation and trophoblast invasion in an already permissive uterine environment. Such memory-like adaptations may contribute to the reduced rates of certain placentation-related complications observed with increasing parity (68,69).

Taken together, these findings support a model of pregnancy as a series of discrete, nonlinear immune transitions that are modulated by fetal and maternal characteristics. Mapping these trajectories at scale provides a framework for distinguishing normative immune adaptation from deviations that may predict pregnancy complications. By identifying which inflammatory mediators exhibit dynamic versus stable patterns, these data help refine candidate biomarker selection for studies of adverse pregnancy outcomes, including hypertensive disorders of pregnancy, gestational diabetes, and spontaneous preterm birth, all of which involve perturbations of the immune–metabolic interface. Furthermore, the observed fetal sex and gravidity effects underscore the importance of incorporating these variables into future analyses rather than treating them as nuisance covariates, as they likely represent biologically meaningful sources of immune variation.

This study has several strengths, including its large and diverse sample drawn from multiple sites within the NIH ECHO Cohort and the breadth of gestational coverage, which spans from 5 to 40 weeks and provides rare insight into early- and late-pregnancy immune states. Nonetheless, some limitations should be acknowledged. The few very early gestational samples (e.g., ∼ 5 weeks) were collected from participants recruited through fertility clinics, who may have been undergoing treatment or medication protocols; these samples may therefore not fully represent immune states in unassisted conceptions. The majority of samples were collected in the third trimester, which could reduce sensitivity for detecting transient early-pregnancy peaks despite the flexibility of GAMs. Relatedly, although effect-modification analyses could be influenced by unequal GA sampling, sensitivity analyses showed no imbalance by fetal sex and only modest trimester differences by gravidity. Early-pregnancy effects are therefore unlikely to reflect sampling bias, though later-trimester stabilization among multigravida participants should be interpreted cautiously. Additionally, although gravidity information was available for the full sample, approximately half of participants self-reported this information (n=621, 53.8%), while the remainder had gravidity extracted from medical records (n=533, 46.2%). Medical record-based gravidity may undercount prior pregnancies, particularly early losses that are less consistently documented. We were also unable to analyze the full panel of 96 proteins included in Olink: proteins with more than 25% of values below the LOD were excluded. Future studies employing more sensitive panels may reveal complementary biology in these low-abundance proteins. Finally, the cross-sectional design precludes direct inference of within-individual change, and longitudinal data will be needed to confirm the timing and magnitude of observed nonlinear transitions.

In summary, this study provides one of the most comprehensive characterizations to date of gestational immune trajectories in a cross-sectional sample of human pregnancies. Maternal inflammation and immune regulation evolve through coordinated, often nonlinear shifts that align with key physiological milestones. These patterns suggest that the maternal immune system operates through flexible, phase-dependent reconfiguration to balance tissue remodeling, tolerance, and metabolic demands across pregnancy. Establishing these normative trajectories provides a critical reference point for identifying early immune deviations that may underlie pregnancy complications and for advancing a more precise understanding of maternal–fetal immunology.

## Author Contributions

S.M.B.: conceptualization, data curation, formal analysis, methodology, visualization, writing – original draft, writing – review & editing; R.S.T.: writing – original draft, writing – review & editing; S.S.: data curation, writing – review & editing; A.G.: conceptualization, funding acquisition, writing – review & editing; C.D.: data curation, writing – review & editing; M.E.N.: data curation, resources, writing – review & editing; L.A.P.: funding acquisition, investigation, writing – review & editing; B.T.: investigation, writing – review & editing; S.M.: writing – review & editing; C.W.H.: funding acquisition, writing – review & editing; A.J.E.: funding acquisition, writing – review & editing; S.N.G.: formal analysis, validation, supervision, writing – original draft, writing – review & editing; L.C.S.: conceptualization, funding acquisition, data curation, methodology, supervision, writing – original draft, writing – review & editing.

## ECHO Acknowledgments

The authors wish to thank our ECHO Colleagues; the medical, nursing, and program staff; and the children and families participating in the ECHO cohort. We would also like to thank Karin Vevang and Hannah Tollman at the HHEAR laboratory for their contributions to the study’s biomarker assays.

## NIH Disclaimer

The content is solely the responsibility of the authors and does not necessarily represent the official views of the National Institutes of Health.

## Data Availability Statement

Select de-identified data from the ECHO Program are available through <u>NICHD’s Data and Specimen Hub (DASH)</u>. Information on study data not available on DASH, such as some Indigenous datasets, can be found on the <u>ECHO study DASH webpage</u>.

