## Supplemental Materials for "A cross-sectional examination of immune adaptations during pregnancy in the ECHO Cohort"

Supplementary Material

Supplementary Table 1. Coefficients of variation. Coefficients of variation (CVs; %) for n=69 proteins included in the analysis are reported in Supplementary Table 1. CVs were calculated within the following groups of samples, which all represented different sets of technical replicates: Control Sample Intra-Plate was calculated within each pair of control samples (technical replicates derived from a pooled plasma sample provided by the data-generating lab) on each plate (n=2 technical replicates per plate; reflects variation in technical replicates derived from a single sample within the same plate); Control Sample Inter-Plate included all control samples (technical replicates derived from a pooled plasma sample provided by the data-generating lab) across all plates (reflects variation in technical replicates derived from a single sample across all plates); Intra-Plate Study Technical Replicate was calculated within each pair of technical replicate sample pairs originating from ECHO biospecimens that were assayed on the same plate within the experiment (reflects variation in technical replicate pairs derived from multiple ECHO biospecimens assayed within the same plate); and Inter-Plate Study Technical Replicate was calculated within each pair of technical replicate samples originating from ECHO biospecimens that were assayed on different plates across the experiment (reflects variation in technical replicates derived from multiple ECHO biospecimens that were assayed across different plates). Abbreviations: CV, coefficient of variation.

| **Protein** | **UniProt ID** | **Olink ID** | **Mean Control Sample Intra-Plate CV** | **Control Sample Inter-Plate CV** | **Mean Intra-Plate Study Technical Replicate CV** | **Mean Inter-Plate Study Technical Replicate CV** |
| --- | --- | --- | --- | --- | --- | --- |
| 4E-BP1 | Q13541 | OID00536 | 16.2 | 31.7 | 11.3 | 17.1 |
| ADA | P00813 | OID00560 | 9.1 | 14.1 | 12.7 | 9.1 |
| AXIN1 | O15169 | OID00487 | 9.9 | 17.7 | 17.8 | 20.7 |
| CASP-8 | Q14790 | OID00550 | 9.7 | 17.7 | 13.9 | 15.6 |
| CCL11 | P51671 | OID00505 | 13.3 | 22.0 | 8.4 | 8.8 |
| CCL19 | Q99731 | OID00513 | 14.8 | 26.1 | 13.9 | 11.1 |
| CCL20 | P78556 | OID00556 | 10.8 | 23.0 | 14.2 | 7.6 |
| CCL23 | P55773 | OID00530 | 14.3 | 27.0 | 12.0 | 10.6 |
| CCL25 | O15444 | OID00551 | 10.1 | 19.2 | 12.1 | 9.1 |
| CCL28 | Q9NRJ3 | OID00539 | 10.1 | 21.1 | 12.0 | 13.9 |
| CCL3 | P10147 | OID00532 | 14.5 | 23.2 | 15.6 | 9.8 |
| CCL4 | P13236 | OID00498 | 14.0 | 24.4 | 11.6 | 9.5 |
| CD244 | Q9BZW8 | OID00477 | 10.2 | 17.1 | 12.4 | 7.9 |
| CD40 | P25942 | OID00542 | 10.5 | 17.5 | 11.4 | 10.6 |
| CD5 | P06127 | OID00531 | 9.3 | 17.8 | 12.7 | 7.8 |
| CD6 | P30203 | OID00499 | 14.3 | 22.5 | 10.3 | 10.5 |
| CD8A | P01732 | OID05124 | 14.9 | 25.4 | 10.8 | 12.1 |
| CDCP1 | Q9H5V8 | OID00476 | 9.5 | 17.7 | 12.1 | 11.2 |
| CSF-1 | P09603 | OID00562 | 6.6 | 9.1 | 5.6 | 6.6 |
| CST5 | P28325 | OID00491 | 10.8 | 16.2 | 7.8 | 7.3 |
| CX3CL1 | P78423 | OID00552 | 11.3 | 17.9 | 14.7 | 12.1 |
| CXCL1 | P09341 | OID00496 | 6.0 | 11.0 | 11.5 | 10.9 |
| CXCL11 | O14625 | OID00486 | 13.1 | 25.1 | 11.5 | 10.3 |
| CXCL5 | P42830 | OID00520 | 12.6 | 20.0 | 16.9 | 12.1 |
| CXCL6 | P80162 | OID00534 | 12.4 | 25.6 | 15.0 | 11.6 |
| CXCL9 | Q07325 | OID00490 | 12.6 | 21.7 | 11.7 | 10.2 |
| DNER | Q8NFT8 | OID01213 | 9.3 | 12.7 | 9.4 | 7.0 |
| EN-RAGE | P80511 | OID00541 | 18.8 | 44.1 | 20.4 | 10.6 |
| FGF-19 | O95750 | OID00545 | 11.3 | 21.4 | 11.8 | 11.5 |
| FGF-21 | Q9NSA1 | OID00512 | 16.9 | 38.9 | 11.9 | 17.2 |
| FGF-23 | Q9GZV9 | OID00507 | 12.5 | 20.0 | 14.3 | 13.5 |
| Flt3L | P49771 | OID00533 | 10.5 | 20.3 | 10.9 | 8.3 |
| HGF | P14210 | OID00522 | 12.8 | 21.9 | 10.5 | 8.7 |
| IFN-γ | P01579 | OID05547 | 11.6 | 22.4 | 10.5 | 11.2 |
| IL10 | P22301 | OID00528 | 11.7 | 22.4 | 12.9 | 8.9 |
| IL-10RB | Q08334 | OID00515 | 12.1 | 14.8 | 7.3 | 9.1 |
| IL-12B | P29460 | OID00523 | 11.3 | 19.7 | 12.4 | 8.0 |
| IL-15RA | Q13261 | OID00514 | 10.5 | 15.7 | 10.6 | 8.8 |
| IL-17C | Q9P0M4 | OID00483 | 12.1 | 25.3 | 10.9 | 16.0 |
| IL18 | Q14116 | OID00501 | 13.9 | 23.4 | 12.6 | 11.6 |
| IL-18R1 | Q13478 | OID00517 | 11.8 | 19.4 | 12.2 | 9.9 |
| IL6 | P05231 | OID00482 | 11.5 | 15.7 | 12.7 | 11.6 |
| IL7 | P13232 | OID00478 | 11.2 | 16.8 | 11.1 | 9.4 |
| IL8 | P10145 | OID00471 | 13.6 | 23.1 | 13.2 | 9.6 |
| LAP-TGF-β-1 | P01137 | OID00480 | 9.9 | 22.1 | 8.7 | 12.1 |
| LIF-R | P42702 | OID00511 | 12.8 | 20.3 | 9.7 | 12.3 |
| MCP-1 | P13500 | OID00484 | 12.6 | 23.5 | 12.8 | 8.0 |
| MCP-2 | P80075 | OID00549 | 10.0 | 21.3 | 13.0 | 10.0 |
| MCP-4 | Q99616 | OID00504 | 13.7 | 25.7 | 11.8 | 8.2 |
| MMP-1 | P03956 | OID00510 | 12.9 | 20.4 | 9.7 | 10.5 |
| MMP-10 | P09238 | OID00527 | 13.4 | 19.4 | 10.5 | 6.9 |
| NT-3 | P20783 | OID00554 | 11.6 | 20.2 | 39.9 | 29.3 |
| OPG | O00300 | OID00479 | 10.6 | 18.1 | 8.8 | 9.2 |
| OSM | P13725 | OID00494 | 10.6 | 20.0 | 9.8 | 11.1 |
| PD-L1 | Q9NZQ7 | OID00518 | 14.9 | 19.2 | 11.4 | 11.1 |
| SCF | P21583 | OID00500 | 9.7 | 11.8 | 5.8 | 8.3 |
| SIRT2 | Q8IXJ6 | OID00538 | 13.4 | 27.0 | 15.8 | 17.5 |
| ST1A1 | P50225 | OID00557 | 8.8 | 18.2 | 16.7 | 20.2 |
| STAMBP | O95630 | OID00558 | 10.5 | 18.5 | 14.9 | 13.8 |
| TGF-α | P01135 | OID00503 | 11.6 | 19.9 | 10.4 | 8.7 |
| TNF | P01375 | OID05548 | 12.3 | 20.4 | 11.9 | 13.1 |
| TNFB | P01374 | OID00561 | 9.0 | 14.5 | 9.9 | 7.8 |
| TNFRSF9 | Q07011 | OID00553 | 8.8 | 15.0 | 11.4 | 9.2 |
| TNFSF14 | O43557 | OID00506 | 13.8 | 21.1 | 10.1 | 7.7 |
| TRAIL | P50591 | OID00488 | 10.4 | 17.6 | 8.3 | 7.9 |
| TRANCE | O14788 | OID00521 | 13.8 | 21.4 | 13.2 | 10.6 |
| TWEAK | O43508 | OID00555 | 12.3 | 24.6 | 14.1 | 9.7 |
| uPA | P00749 | OID00481 | 10.6 | 20.6 | 7.8 | 9.7 |
| VEGFA | P15692 | OID00472 | 11.9 | 21.1 | 12.3 | 8.0 |

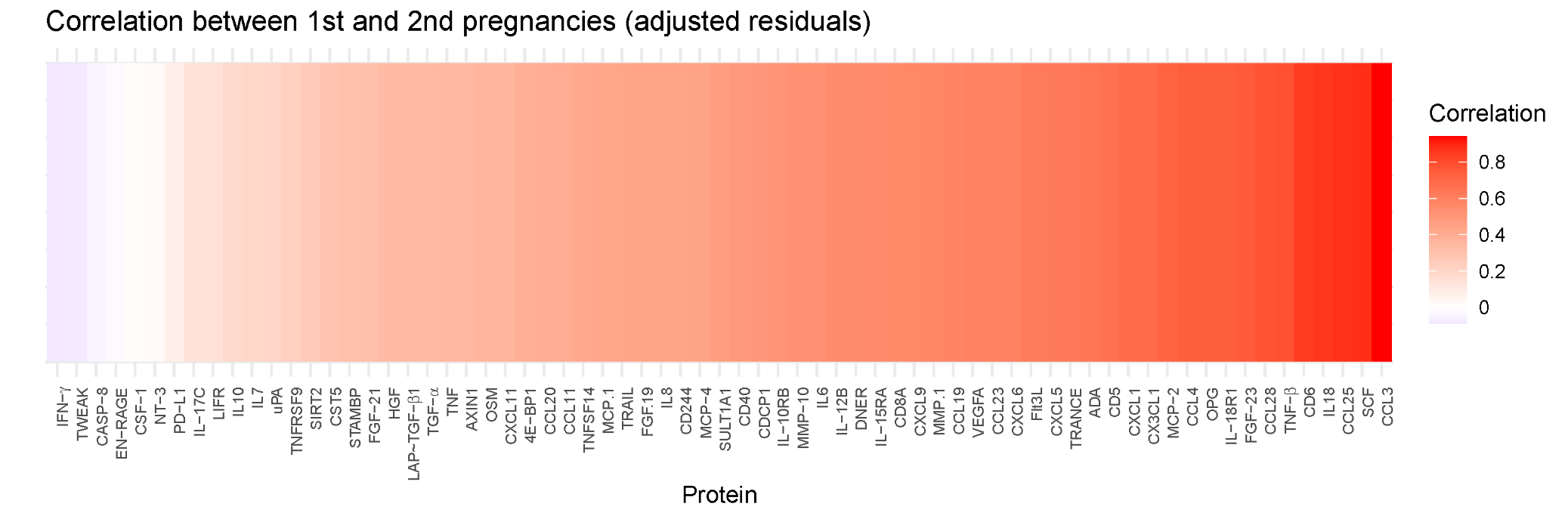

**Supplementary Figure 1.** **Within-subject correlations in protein expression levels between subsequent pregnancies.** After regressing out the effects of gestational age at blood draw, maternal age, fetal sex, and gravidity, the majority of proteins showed consistent expression levels across pregnancies within a participant. Only 4 of the 69 proteins showed correlation coefficients < 0: IFN-γ, TWEAK, CASP-8, and EN-RAGE. To evaluate within-person consistency of inflammatory protein expression across pregnancies, we identified participants with at least two pregnancies (n=30). Participants missing any protein or covariate value (gestational age at blood draw, maternal age, fetal sex, and gravidity) in either pregnancy were excluded (n=12, resulting in n=18 to be analyzed). For each participant, pregnancies were chronologically ordered based on blood draw date, and only the first two pregnancies were retained. We additionally excluded pregnancies with implausible gravidity patterns (e.g., gravidity not increasing across pregnancies). Protein values were residualized to remove effects of gestational age, maternal age, fetal sex, and gravidity using linear regression models. For each protein, residual values from pregnancy 1 and pregnancy 2 were reshaped into a wide format and pairwise Pearson correlations were computed.

Supplementary Table 2. Functional classification of the 69 inflammatory proteins from the Olink Target 96 Inflammation panel included in analyses. A list of gene names, alternative names, and biomarker names (all as defined by Olink) is provided for the 69 proteins included in our analyses. For each protein, we additionally provide a description of its general biological function based on definitions from the National Institutes of Health gene database (1) and a summary of its known or proposed roles in pregnancy-related physiological processes and/or pregnancy-associated conditions.

| **Gene Name** | **Alternative Names** | **Biomarker** | **Role in Pregnancy** |
| --- | --- | --- | --- |
| CCL2 | MCP1, SCYA2 | C-C motif chemokine 2 | Cytokine that mediates the recruitment of monocytes. Expressed by first-trimester decidual stromal cells (DSCs), promoting primary DSC proliferation and growth (2). Abnormal levels associated with adverse pregnancy outcomes (3). Increased in gestational diabetes mellitus (GMD) (4). Associated with anxiety in pregnancy (5). |
| CCL3 | MIP1A, G0S19, SCYA3 | C-C motif chemokine 3 | Chemokine involved in adaptive immune response and inducible in hematopoietic cells. Increases rapidly in the first trimester when trophoblasts differentiate to extravillous trophoblasts (2). |
| CCL4 | MIP1B, LAG1, SCYA4 | C-C motif chemokine 4 | Chemokine involved in adaptive immune response and inducible in hematopoietic cells. Involved in uterine receptivity and placentation in early pregnancy (6). Decreased in GDM (4). Associated with anxiety and depression in pregnancy (5). |
| CCL8 | MCP2, SCYA10, SCYA8 | C-C motif chemokine 8 | Inflammatory chemokine expressed by fibroblasts and associated with Th2 immune response. In early pregnancy, CCL8 from DSCs supports proper macrophage positioning and immune tolerance. Upregulated in hypoxic states. High levels contribute to the pathogenesis of preeclampsia (7). |
| CCL11 | SCYA11 | Eotaxin | Eosinophil chemoattractant implicated in neuroinflammation and neurodegenerative disorders (8). In pregnancy, plays a pro-invasive role in decidual control of trophoblast invasion (9). Associated with depression in pregnancy (5). |
| CCL13 | MCP4, NCC1, SCYA13 | C-C motif chemokine 13 | Chemotactic agent for many immune cells with an established role in rheumatologic, dermatologic, and oncologic disease. In pregnant patients with multiple sclerosis, decreased in late pregnancy and increased in the postpartum period (10). |
| CCL19 | MIP3B, ELC, CYA19 | C-C motif chemokine 19 | Chemokine expressed at high levels in the thymus and lymph nodes that directs a variety of cell types to T-cell zones of lymphoid organs. Physiologically decreased in pregnancy with thymic involution (11). |
| CCL20 | MIP3A, LARC, SCYA20 | C-C motif chemokine 20 | Upregulated in the plasma of patients with preeclampsia in early and late pregnancy (12). Increased levels in the lower segment myometrium found in preterm labor (13). |
| CCL23 | MPIF1, MIP3, SCYA23 | C-C motif chemokine 23 | Chemoattractant of T lymphocytes, monocytes and neutrophils and angiogenesis mediator. In preeclampsia, upregulated in those with small-for-gestational age infants (14). |
| CCL25 | TECK, SCYA25 | C-C motif chemokine 25 | Chemokine highly expressed by thymus and intestinal epithelium with role in mucosal immunity. Limited literature on role in pregnancy. |
| CCL28 | SCYA28 | C-C motif chemokine 28 | Chemoattractant of CD4 or CD8 T cells and eosinophils involved in the mucosal homing of B and T cells. Physiologically decreased in pregnancy with thymic involution (11). |
| CXCL1 | GRO, GRO1, GROA, MGSA, SCYB1 | Growth-regulated alpha protein | Monocyte chemoattractant secreted by activated T cells and implicated in various inflammatory diseases. May stimulate angiogenesis in the first trimester via the vascular endothelial growth factor pathway (15). Increased in GDM (4). |
| CXCL5 | ENA78, SCYB5 | C-X-C motif chemokine 5 | Neutrophil chemoattractant and promoter of angiogenesis. Abnormal secretion of CXL5 may be early marker of spontaneous pregnancy loss; downregulated in the villous tissue of patients with recurrent spontaneous pregnancy loss (2). |
| CXCL6 | GCP2, SCYB6 | C-X-C motif chemokine 6 | Proinflammatory chemokine implicated in connective tissue metabolism and neuroinflammatory disorders. In pregnancy, inhibits trophoblast cell migration and invasion in the first trimester (16). Increased expression in perinatal mood and anxiety disorders in patients without preeclampsia (17). |
| CXCL9 | MIG, CMK, SCYB9 | C-X-C motif chemokine 9 | Activated T lymphocyte chemoattractant. Increased in GDM (4). Increased in the second trimester in pregnancies resulting in preterm birth (18). |
| CXCL11 | ITAC, SCYB11, SCYB9B | C-X-C motif chemokine 11 | Activated T lymphocyte chemoattractant. Excess CXCL11 in decidua blunts peripheral natural killer (NK) cell recruitment and may contribute to shallow placentation in preeclampsia (2). Decreased in GDM (4). Increased expression in perinatal mood and anxiety disorders in patients without preeclampsia (17). |
| CX3CL1 | FKN, NTT, SCYD1 | Fractalkine | Monocyte chemoattractant with role in wide range of inflammatory, oncologic, and infectious disease. Highly expressed by cytotrophoblast decidual cells. Elevated levels associated with preterm labor and a predictive biomarker of preterm premature rupture of membranes (19). Also associated with early pregnancy loss. Elevated levels in pregnancy associated with increased maternal lifetime stress and trauma (20). |
| IL6 | IFNB2 | Interleukin-6 | Cytokine with major role in inflammation and the maturation of B cells and an endogenous pyrogen, widely implicated in disease states. Proposed role in implantation and remodeling of the spiral arteries in physiologic pregnancy; abnormal expression implicated in pregnancy loss, preeclampsia, and GDM (21). |
| IL8 | CXCL8 | Interleukin-8 | Proinflammatory cytokine with major role in mediating inflammatory response. Neutrophil chemoattractant and potent angiogenic factor. Implicated in physiologic pregnancy establishment and maintenance; abnormal expression implicated in pregnancy loss, preeclampsia, and GDM (21). Elevated levels associated with increased anxiety in pregnancy (5). |
| IL12B | NKSF2 | Interleukin-12 subunit beta | Cytokine expressed by activated macrophages that induces Th1 cell development. Polymorphisms of this gene associated with preeclampsia (22). |
| IL17C | None | Interleukin-17C | Cytokine expressed by activated T cells that stimulates the release of TNF-alpha and IL1-beta from monocytes. Increased in preeclamptic pregnancies resulting in small-for-gestational-age infants (14). Otherwise, limited literature on role in pregnancy. |
| IL18 | IGIF, IL1F4 | Interleukin-18 | Proinflammatory cytokine implicated in the injury of various organs and in cytokine storms. Limited literature on role in pregnancy. |
| TNF | TNFA, TNFSF2 | Tumor necrosis factor | Multifunctional proinflammatory cytokine primarily secreted by macrophages. Involved in the regulation of many biologic processes and implicated in various immunologic diseases. TNF-alpha influences hormone synthesis, placental architecture, and embryonic development in physiologic pregnancy, and increased levels are associated with poor pregnancy outcomes including early pregnancy loss and preeclampsia (23). |
| TNFSF10 | TRAIL, APO2L | Tumor necrosis factor ligand superfamily member 10 | Cytokine that preferentially induces apoptosis in transformed and tumor cells. Expressed at the maternal–fetal interface to induce apoptosis of lymphocytes to maintain immune tolerance. Induces apoptosis of vascular smooth muscle cells for spiral artery remodeling in physiologic pregnancy. Decreased levels in preeclampsia (24). |
| TNFSF12 | TWEAK, APO3L, DRLG | Tumor necrosis factor ligand superfamily member 12 | Cytokine that induces apoptosis via multiple pathways and regulates angiogenesis. Expressed in the placenta may contribute to cell death and immune regulation at the maternal–fetal interface. Low levels associated with preeclampsia and GDM (25,26). |
| TNFSF14 | LIGHT, HVEML | Tumor necrosis factor ligand superfamily member 14 | Herpesvirus entry mediator that also stimulates T cell proliferation and apoptosis. Expressed in the placenta and may contribute to cell death and immune regulation at the maternal–fetal interface. Elevated in preeclampsia (27). |
| IFNG | None | Interferon gamma | Cytokine secreted by innate and adaptive immune system cells that triggers cellular response to viral and microbial infections. In physiologic pregnancy, mediates uterine spiral artery remodeling; aberrant expression leads to embryotoxicity and abnormal placental development (28). |
| S100A12 | EN-RAGE | Protein S100-A12 | Calcium-binding protein that regulates cellular processes including cell cycle progression and may mediate neutrophil function. Increased levels in preeclampsia, chorioamnioniitis, and polycystic ovary syndrome (29). |
| IL10 | None | Interleukin-10 | Cytokine with many effects on inflammation and immunoregulation. Plays a major anti-inflammatory and immunoregulatory role in pregnancy, maintaining immune tolerance. Decreased levels associated with adverse pregnancy complications (preterm birth, miscarriage, fetal growth restriction) and preeclampsia (30). |
| TGFB1 (LAP form) | TGFB | Transforming growth factor-β1 proprotein | Pleiotropic cytokine that regulates cell proliferation and growth; predominantly anti-inflammatory in effect and frequently elevated in tumor cells. As immunosuppressive, regulates several aspects of pregnancy, such as trophoblast uterine invasiveness. Elevated in preeclampsia (31). Altered expression may contribute to the pathogenesis of recurrent pregnancy loss (32). |
| LTA | TNFB, TNFSF1 | Lymphotoxin-alpha | Cytokine produced by lymphocytes as part of tumor necrosis factor protein family. Proinflammatory and pro-apoptotic; mediates various inflammatory, immunostimulatory, and antiviral responses. Implicated in preeclampsia and pregnancy loss, with polymorphisms associated with recurrent pregnancy loss (33). |
| VEGFA | VEGF | Vascular endothelial growth factor A | Induces proliferation and migration of vascular endothelial cells. Essential for physiologic and pathologic angiogenesis. In pregnancy, supports placental formation, spiral artery remodeling, and embryo implantation and plays a role in immune regulation. Dysregulated in preeclampsia, early pregnancy loss, recurrent miscarriage, and fetal growth restriction (34). |
| HGF | HGF | Hepatocyte growth factor | Pleiotropic factor that regulates cell growth, motility, morphogenesis, and repair in various cell types. In pregnancy, stimulates trophoblast invasion and contributes to placental vascular development as well as fetal organ development. Reduced in preeclampsia and intrauterine growth restriction (35). |
| FGF19 | None | Fibroblast growth factor 19 | Metabolic hormone with significant role in bile acid metabolism and growth factor involved in embryonic nervous and skeletal system development (36). Reduced levels in insulin resistance, including GDM. Positively correlated with birth weight in female infants (37). |
| FGF21 | None | Fibroblast growth factor 21 | Metabolic hormone that promotes insulin sensitivity. Rises in the third trimester as metabolic demands increase. Elevated in GDM and preeclampsia, likely to compensate for insulin resistance (38). |
| FGF23 | HYPF | Fibroblast growth factor 23 | Bone-derived protein that regulates renal phosphate and vitamin D homeostasis. Plays a role in the pathogenesis of gestational hypertension and fetal growth disorders; potential biomarker for fetal growth (39). |
| KITLG | Stem cell factor (SCF), MGF | Kit ligand | Pleiotropic growth factor involved in stem cell differentiation and fetal germ cell development, neural cell development, and hematopoiesis. Expressed by the placental villous core and may regulate trophoblast proliferation (40). Activates primordial follicles and plays role in fertility (41). |
| CSF1 | None | Macrophage colony-stimulating factor (M-CSF) 1 | Cytokine involved in macrophage production, differentiation, and function. In pregnancy, regulates placental development function. Dysregulation may contribute to preeclampsia and gestational hypertension (42). |
| LIFR | None | Leukemia inhibitory factor receptor | Receptor for leukemia inhibitory factor, a cytokine in the IL-6 family involved in fetal and adult cellular differentiation and proliferation. Regulates endometrial receptivity and implantation, including trophoblast cells and macrophage responses to inflammatory cytokines (43). |
| OSM | None | Oncostatin-M | IL-6 family cytokine and growth regulator that inhibits tumor cell line proliferation and regulates the production of various cytokines. Contributes to embryo implantation and decidualization and stimulates release of human chorionic gonadotrophin (44). |
| TGFA | None | Protransforming growth factor-α | Growth factor and ligand for epidermal growth factor receptor associated with multiple cancers. In physiologic pregnancy, serves as a major anti-inflammatory cytokine and stimulates placental mitosis. Elevated in cord blood of neonates with intrauterine growth restriction (45). |
| NTF3 | None | Neurotrophin-3 | Neurotrophin that may influence embryonic neural development and contribute to angiogenesis during embryogenesis; not significantly associated with fetal growth disorders (46). |
| PLAU | None | Urokinase-type plasminogen activator (uPA) | Serine protease that converts plasminogen to plasmin, a key step in fibrinolysis. Regulates trophoblast invasion and migration in early placentation (47). |
| MMP1 | CLG | Interstitial collagenase | Protease that breaks down interstitial collagens (type I and III). A physiologic constituent of amniotic fluid that increases with gestational age; increased in premature preterm rupture of membranes (48). |
| MMP10 | STMY2 | Stromelysin-2 | Protease that breaks down various extracellular matrix proteins including collagens, fibronectin, elastin, and laminin. Widely expressed at the maternal–fetal interface in pregnancy and stimulated by interleukin 1β (49). Reduced levels in preeclampsia (50). |
| TNFSF11 | TRANCE, RANKL, OPGL | Tumor necrosis factor ligand superfamily member 11 | Ligand for osteoprotegerin with major role in osteoclast differentiation and activation that also serves as a dendritic cell survival factor involved in regulating cell-mediated immunity. Elevated levels observed postpartum (51). Polymorphisms associated with preeclampsia (52). |
| TNFRSF11B | OPG, OCIF | Tumor necrosis factor receptor superfamily member 11B | Osteoprotegerin ligand and decoy receptor secreted by osteoblasts that downregulates bone reabsorption. May contribute to the regulation of bone turnover in pregnancy, but the literature is limited (53). |
| CD274 | B7H1, PDCD1L1, PDCD1LG1, PDL1 | Programmed cell death 1 ligand 1 | Immune inhibitory receptor ligand that inhibits T-cell activation and cytokine production. Physiologically, prevents the development of autoimmunity; in tumors, can allow immune evasion. In pregnancy, expressed by the placenta, especially by syncytiotrophoblasts and cytotrophoblasts, to maintain immune tolerance, preventing fetal rejection (54). Per animal models, signaling of this pathway may be altered in pregnancy loss (55). Expression is decreased in preterm labor (56). |
| CD244 | 2B4 | NK cell receptor 2B4 | Cell surface receptor primarily expressed by NK cells that mediates cytotoxicity outside of the major histocompatibility complex. In pregnancy, expression in decidual NK cells contributes to immunoregulation and fetal tolerance (57). Reduced expression found in recurrent pregnancy loss (58). |
| CD40 | TNFRSF5 | Tumor necrosis factor receptor superfamily member 5 | Receptor on antigen-presenting cells that serves as a key immune checkpoint, mediating T cell-dependent immunoglobulin class switching, memory B cell development, and germinal center formation. Per animal models, may play a role in the establishment of pregnancy through regulating endometrial endothelial cell function during implantation (59). Its ligand, soluble CL40L, is increased in preeclampsia and other hypertensive disorders of pregnancy (60). |
| CD5 | LEU1 | T-cell surface glycoprotein CD5 | T cell, B1-a cell, and B cell chronic lymphocytic leukemia cell receptor that modulates T and B cell signaling. CD5+CD1d+ B regulatory cells are decreased in healthy pregnancy (61), while CD19+ CD5+ B cells are reduced in preeclampsia (62); however, literature on the receptor's specific role in pregnancy is limited. |
| CD6 | None | T-cell differentiation antigen CD6 | T cell membrane protein that contributes to T cell activation. Limited literature in pregnancy. |
| CD8A | MAL | T-cell surface glycoprotein CD8 alpha chain | Receptor found on most cytotoxic T cells that acts as a co-receptor for antigens displayed by antigen-presenting cells with major histocompatibility complex class 1 molecules. Used as a marker for CD8+ T-cells, which are abundant at the maternal–fetal interface; however, the literature on the receptor's specific role in pregnancy is limited (63). |
| CDCP1 | TRASK | CUB domain-containing protein 1 | Transmembrane protein with role in regulating cellular events during tumor invasion and metastasis. Expressed by a subset of trophoblasts, but literature is limited in pregnancy (64). |
| IL10RA | IL10R | Interleukin-10 receptor subunit alpha | Protein that mediates interleukin 10 signaling and inhibits proinflammatory cytokines. Coexpression with IL10RB required for IL-10 signaling. IL-10R is increased in the early pregnant endometrium and may act as an anti-inflammatory paracrine mediator at the maternal-fetal interface (65). |
| IL10RB | CRFB4, D21S58, D21S66 | Interleukin-10 receptor subunit beta | See above; coexpression with IL10RA required for IL-10 signaling. |
| IL15RA | None | Interleukin-15 receptor subunit alpha | Cytokine receptor that binds interleukin 15 with high affinity and specificity. IL15RA is expressed by the placenta and by decidual NK cells; IL-15 stimulates in vitro proliferation of decidual NK cells without transforming them to be cytotoxic to trophoblasts (66). IL-15 is increased in preeclampsia and decreased in implantation failure (67). |
| IL18R1 | IL1RRP | Interleukin-18 receptor 1 | Cytokine receptor in the interleukin 1 receptor family that specifically binds interleukin 18 and is expressed by the endometrium and by villous trophoblasts. In early pregnancy, IL18 may regulate the cytotoxicity of NK cells. IL18 is also elevated during labor and in various pregnancy complications (68). |
| TNFRSF9 | CD137, ILA | Tumor necrosis factor receptor superfamily member 9 | Receptor in the TNF-receptor family that contributes to T cell development, survival, and clonal expansion as well as proliferation of peripheral monocytes. In pregnancy, expression increases in the uterus during implantation in animal models, but its function is unknown (69). |
| AXIN1 | AXIN | Axis inhibition protein 1 | Cytoplasmic protein that can induce apoptosis as an antagonist of the WNT/β-catenin signaling pathway; mutations associated with various cancers. Higher expression found in intrauterine growth restriction placentae (70). Lower levels in late pregnancy associated with postpartum depression (51). |
| CST5 | None | Cystatin D | Type 2 cystatin with expression restricted to the submandibular and parotid glands (71). No known role in pregnancy. |
| DNER | BET | Delta and Notch-like EGF-related receptor | Protein expressed by the developing and mature central nervous system (72). No known role in pregnancy. |
| EIF4EBP1 | 4E-BP1 | Eukaryotic translation initiation factor 4E-binding protein 1 | Translation repressor protein that binds to eukaryotic translation initiation factor 4E (eIF4E) to inhibit protein translation. In animal models, plays a role in spindle formation in meiosis and in the transition of maternal to embryonic control of development (73). Otherwise, literature is limited in pregnancy. |
| SIRT2 | SIR2L, SIR2L2 | NAD-dependent protein deacetylase sirtuin-2 | Protein involved in carbohydrate metabolism and the cell cycle with emerging role in female fertility (74). Found in placental syncytiotrophoblasts and downregulated in preeclampsia (75). |
| STAMBP | AMSH | STAM-binding protein | Deubiquitinating enzyme with a defective form leading to microcephaly-capillary malformation syndrome. Lower levels in late pregnancy associated with postpartum depression (76). |
| SULT1A1 | STP, STP1 | Sulfotransferase 1A1 | Sulfate conjugation enzyme functional in the placenta, with substrates that include endogenous hormones (estradiol, thyroid hormones) and endogenous substances such as medicinal drugs and environmental chemicals, with likely role in mediating fetal xenobiotic exposure (77). Lower levels in late pregnancy associated with postpartum depression (76). |
| CASP8 | MCH5 | Caspase-8 | Apoptotic initiation factor involved in Fas-induced programmed cell death and other apoptotic pathways. Low expression in physiologic pregnancy but elevated in preterm labor, preeclampsia, and intrauterine growth restriction (78). |
| ADA | ADA1 | Adenosine deaminase | Enzyme in the purine catabolic pathway essential for lymphoid cell differentiation, defective in severe combined immunodeficiency disease. Increased in recurrent pregnancy loss, preeclampsia, and hyperemesis gravidarum (79). Lower levels in late pregnancy associated with postpartum depression (76). |

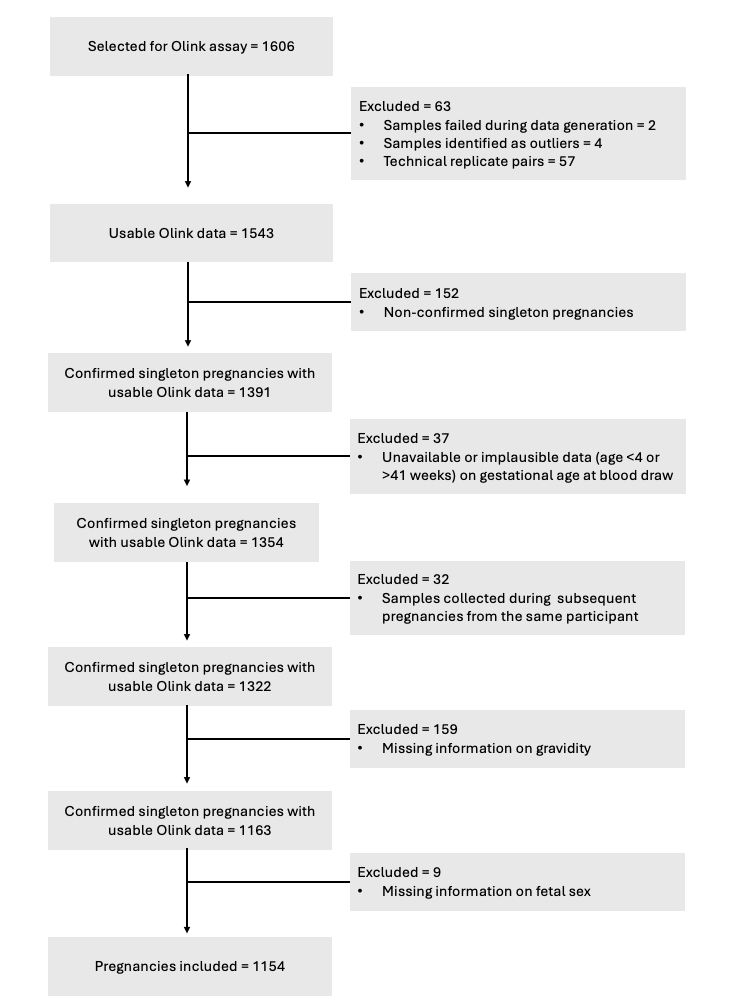

**Supplementary Figure 2. Flow chart depicting each stage of participant exclusions.** A total of 1154 pregnancies were included in the final analysis.

Supplementary Table 3. Trimester differences in protein expression levels. To identify markers that differed by trimester, we fit separate linear mixed-effects regression models for each protein, predicting expression level from trimester. Models controlled for maternal age at blood draw, gravidity, and infant sex and contained a random intercept for site to account for site-level heterogeneity. Because sample sizes differed across trimesters, we used linear mixed-effects regression with restricted maximum likelihood estimation, which provides valid estimates under unbalanced designs. Overall trimester effects were evaluated using Type III sums of squares, yielding an omnibus F-test for the null hypothesis that mean expression levels were equal across trimesters. Multiple testing was addressed by controlling the false discovery rate (FDR) at 5% across all proteins using the Benjamini–Hochberg procedure. To visualize the direction of trimester effects, we extracted the fixed-effect coefficients for the third trimester contrasts (relative to the first trimester) and ordered proteins by the sign and magnitude of these estimates. When modeling trimester as a categorical predictor, we observed widespread differences in protein expression across pregnancy. Mixed-effects regression models identified 53 out of 69 inflammatory proteins that showed significant differences after correcting for multiple comparisons with an FDR of 5% (Supplementary Table 3; Supplementary Figure 3). These trimester-related changes spanned multiple functional groups, including chemokines (e.g., CCL28, CCL11), growth factors (e.g., LIFR, uPA), pro-inflammatory cytokines (e.g., TWEAK), and other regulatory or signaling proteins (e.g., PD-L1, CD8A). Abbreviations: AIC, Akaike information criterion; CI, confidence interval; FDR, false discovery rate.

| **Protein** | **T1–T2 β estimate** | **T1–T2 t statistic** | **T1–T2 p** | **T1–T2 95% CI** | **T1–T3 B estimate** | **T1–T3 t statistic** | **T1–T3 p** | **T1–T3 95% CI** | **AIC** | **Omnibus F** | **Omnibus p (uncorrected)** | **Omnibus p (FDR-adjusted)** |
| --- | --- | --- | --- | --- | --- | --- | --- | --- | --- | --- | --- | --- |
| PD-L1 | 1.129 | 17.554 | 2.84E-61 | [1.003, 1.255] | 1.696 | 25.396 | 8.55E-111 | [1.565, 1.827] | 1767.670 | 322.839 | 0.000 | 4.73E-109 |
| LIF-R | 1.018 | 15.161 | 1.86E-47 | [0.887, 1.150] | 1.478 | 21.383 | 1.01E-80 | [1.342, 1.613] | 1866.333 | 229.301 | 0.000 | 1.50E-80 |
| TWEAK | -0.653 | -14.216 | 2.30E-42 | [-0.743, -0.563] | -0.750 | -15.919 | 2.23E-49 | [-0.842, -0.658] | 994.186 | 138.191 | 0.000 | 8.19E-52 |
| uPA | 0.168 | 3.528 | 4.36E-04 | [0.074, 0.261] | 0.656 | 13.213 | 3.66E-37 | [0.559, 0.754] | 1076.232 | 111.976 | 0.000 | 6.85E-44 |
| CDCP1 | 0.289 | 4.009 | 6.50E-05 | [0.148, 0.430] | 0.688 | 11.478 | 6.02E-29 | [0.571, 0.806] | 2039.151 | 89.721 | 0.000 | 8.67E-36 |
| CCL28 | 1.200 | 10.453 | 1.76E-24 | [0.975, 1.425] | 1.937 | 18.530 | 4.25E-31 | [1.732, 2.142] | 3101.498 | 178.804 | 0.000 | 1.21E-32 |
| LAP-TGF-β-1 | 0.093 | 1.463 | 1.44E-01 | [-0.032, 0.219] | 0.666 | 9.989 | 1.45E-22 | [0.536, 0.797] | 1753.690 | 73.885 | 0.000 | 6.71E-30 |
| IL-10RB | 0.222 | 6.577 | 7.26E-11 | [0.156, 0.288] | 0.403 | 11.643 | 6.05E-29 | [0.335, 0.471] | 284.321 | 69.107 | 0.000 | 9.27E-28 |
| CD8A | -0.693 | -7.163 | 1.41E-12 | [-0.882, -0.503] | -1.135 | -11.387 | 4.28E-28 | [-1.331, -0.940] | 2702.830 | 65.014 | 0.000 | 2.21E-26 |
| TRANCE | -0.504 | -7.160 | 1.44E-12 | [-0.642, -0.366] | -0.773 | -11.002 | 2.60E-24 | [-0.911, -0.635] | 1972.979 | 60.600 | 0.000 | 1.79E-23 |
| CCL11 | -0.531 | -8.857 | 3.08E-18 | [-0.648, -0.413] | -0.627 | -10.629 | 2.32E-21 | [-0.742, -0.511] | 1604.189 | 58.927 | 0.000 | 1.03E-21 |
| MCP-4 | -0.721 | -7.553 | 8.68E-14 | [-0.908, -0.534] | -0.951 | -9.588 | 6.39E-21 | [-1.146, -0.757] | 2674.359 | 47.192 | 0.000 | 1.25E-19 |
| IL-12B | -0.605 | -8.070 | 1.77E-15 | [-0.752, -0.458] | -0.673 | -8.901 | 1.46E-17 | [-0.822, -0.525] | 2116.587 | 43.546 | 0.000 | 1.04E-17 |
| OPG | 0.260 | 3.563 | 3.82E-04 | [0.117, 0.403] | 0.632 | 8.546 | 1.77E-16 | [0.487, 0.777] | 2055.478 | 40.879 | 0.000 | 8.88E-17 |
| DNER | -0.238 | -6.977 | 5.08E-12 | [-0.305, -0.171] | -0.264 | -7.636 | 1.16E-13 | [-0.331, -0.196] | 312.901 | 32.237 | 0.000 | 1.92E-13 |
| FGF-23 | 0.422 | 2.158 | 3.12E-02 | [0.039, 0.805] | 1.402 | 7.325 | 1.02E-11 | [1.027, 1.778] | 4318.647 | 34.404 | 0.000 | 2.59E-13 |
| CCL23 | -0.390 | -6.077 | 1.67E-09 | [-0.515, -0.264] | -0.489 | -7.387 | 3.48E-13 | [-0.618, -0.359] | 1759.821 | 28.464 | 0.000 | 3.83E-12 |
| MCP-2 | -0.635 | -6.769 | 2.07E-11 | [-0.819, -0.451] | -0.663 | -6.987 | 8.74E-12 | [-0.849, -0.477] | 2631.874 | 28.308 | 0.000 | 5.74E-12 |
| TNFSF14 | -0.465 | -6.233 | 6.41E-10 | [-0.612, -0.319] | -0.530 | -6.814 | 1.56E-11 | [-0.683, -0.378] | 2111.311 | 25.812 | 0.000 | 3.98E-11 |
| CD5 | -0.276 | -6.003 | 2.60E-09 | [-0.367, -0.186] | -0.332 | -6.911 | 8.08E-12 | [-0.427, -0.238] | 1002.979 | 25.659 | 0.000 | 4.35E-11 |
| SCF | -0.302 | -5.945 | 3.67E-09 | [-0.402, -0.202] | -0.346 | -7.479 | 2.51E-11 | [-0.436, -0.255] | 1230.348 | 28.390 | 0.000 | 2.25E-10 |
| CSF-1 | 0.149 | 5.816 | 7.79E-09 | [0.099, 0.199] | 0.172 | 6.498 | 1.36E-10 | [0.120, 0.224] | -344.127 | 23.083 | 0.000 | 4.96E-10 |
| VEGFA | 0.156 | 4.598 | 4.74E-06 | [0.089, 0.222] | 0.225 | 6.389 | 2.51E-10 | [0.156, 0.294] | 296.896 | 20.513 | 0.000 | 5.37E-09 |
| TNFRSF9 | -0.295 | -6.039 | 2.09E-09 | [-0.391, -0.199] | -0.269 | -5.438 | 8.55E-08 | [-0.366, -0.172] | 1135.637 | 19.959 | 0.000 | 1.09E-08 |
| CXCL5 | -0.564 | -2.931 | 3.44E-03 | [-0.942, -0.187] | -1.183 | -5.872 | 5.65E-09 | [-1.577, -0.788] | 4286.667 | 18.083 | 0.000 | 5.12E-08 |
| EN-RAGE | -0.399 | -4.633 | 4.01E-06 | [-0.568, -0.230] | -0.509 | -5.661 | 1.92E-08 | [-0.685, -0.333] | 2440.568 | 16.700 | 0.000 | 1.89E-07 |
| CXCL6 | -0.479 | -4.457 | 9.12E-06 | [-0.689, -0.268] | -0.625 | -5.581 | 3.01E-08 | [-0.845, -0.406] | 2946.305 | 16.073 | 0.000 | 3.35E-07 |
| CX3CL1 | -0.270 | -4.888 | 1.16E-06 | [-0.378, -0.162] | -0.298 | -5.338 | 1.45E-07 | [-0.407, -0.188] | 1413.920 | 15.780 | 0.000 | 4.97E-07 |
| TRAIL | -0.209 | -5.343 | 1.10E-07 | [-0.285, -0.132] | -0.192 | -4.841 | 1.69E-06 | [-0.270, -0.114] | 621.619 | 15.698 | 0.000 | 5.01E-07 |
| CD40 | -0.300 | -5.330 | 1.18E-07 | [-0.410, -0.189] | -0.266 | -4.534 | 6.40E-06 | [-0.382, -0.151] | 1461.347 | 15.087 | 0.000 | 7.85E-07 |
| CD244 | -0.230 | -5.021 | 5.96E-07 | [-0.320, -0.140] | -0.228 | -4.895 | 1.27E-06 | [-0.320, -0.137] | 988.672 | 14.721 | 0.000 | 1.18E-06 |
| STAMBP | -0.645 | -5.213 | 2.20E-07 | [-0.888, -0.403] | -0.510 | -3.956 | 8.12E-05 | [-0.763, -0.257] | 3270.799 | 13.803 | 0.000 | 2.58E-06 |
| IL7 | -0.294 | -4.195 | 2.94E-05 | [-0.431, -0.157] | -0.367 | -5.008 | 6.38E-07 | [-0.510, -0.223] | 1966.895 | 13.212 | 0.000 | 4.45E-06 |
| CD6 | -0.399 | -4.937 | 9.11E-07 | [-0.558, -0.241] | -0.332 | -3.912 | 9.70E-05 | [-0.499, -0.166] | 2302.094 | 12.549 | 0.000 | 8.25E-06 |
| IL18 | 0.231 | 3.126 | 1.81E-03 | [0.086, 0.377] | 0.370 | 4.974 | 9.99E-07 | [0.224, 0.515] | 2088.916 | 12.418 | 0.000 | 1.05E-05 |
| IL-15RA | 0.046 | 1.229 | 2.19E-01 | [-0.028, 0.121] | 0.160 | 4.160 | 3.70E-05 | [0.084, 0.235] | 547.796 | 10.902 | 0.000 | 4.15E-05 |
| SIRT2 | -0.712 | -4.504 | 7.36E-06 | [-1.022, -0.402] | -0.636 | -3.849 | 1.25E-04 | [-0.960, -0.312] | 3834.757 | 10.797 | 0.000 | 4.22E-05 |
| HGF | -0.016 | -0.301 | 7.64E-01 | [-0.122, 0.090] | 0.169 | 3.050 | 2.39E-03 | [0.060, 0.277] | 1372.932 | 9.652 | 0.000 | 1.31E-04 |
| NT-3 | -0.242 | -3.808 | 1.48E-04 | [-0.367, -0.117] | -0.260 | -4.363 | 2.82E-05 | [-0.376, -0.143] | 1742.303 | 10.087 | 0.000 | 1.35E-04 |
| MCP-1 | -0.207 | -3.306 | 9.74E-04 | [-0.330, -0.084] | -0.259 | -4.495 | 1.97E-05 | [-0.371, -0.146] | 1713.553 | 10.122 | 0.000 | 1.51E-04 |
| MMP-10 | -0.042 | -0.555 | 5.79E-01 | [-0.188, 0.105] | -0.197 | -3.166 | 1.59E-03 | [-0.319, -0.075] | 2126.917 | 8.749 | 0.000 | 2.85E-04 |
| AXIN1 | -0.614 | -4.090 | 4.62E-05 | [-0.909, -0.320] | -0.508 | -3.222 | 1.31E-03 | [-0.817, -0.199] | 3720.048 | 8.593 | 0.000 | 3.25E-04 |
| IFN-γ | -0.283 | -2.058 | 3.98E-02 | [-0.553, -0.014] | -0.515 | -4.050 | 1.13E-04 | [-0.765, -0.266] | 3514.270 | 8.806 | 0.000 | 4.54E-04 |
| CCL19 | -0.353 | -3.833 | 1.33E-04 | [-0.533, -0.172] | -0.323 | -3.561 | 4.50E-04 | [-0.501, -0.145] | 2590.156 | 8.190 | 0.000 | 5.26E-04 |
| IL-17C | 0.161 | 1.886 | 5.96E-02 | [-0.006, 0.328] | 0.338 | 3.845 | 1.29E-04 | [0.166, 0.510] | 2411.060 | 7.812 | 0.000 | 6.59E-04 |
| IL8 | -0.231 | -2.655 | 8.05E-03 | [-0.402, -0.060] | -0.349 | -3.876 | 1.14E-04 | [-0.525, -0.172] | 2461.684 | 7.516 | 0.001 | 8.62E-04 |
| TNFB | -0.176 | -3.302 | 9.89E-04 | [-0.281, -0.072] | -0.183 | -3.592 | 4.34E-04 | [-0.284, -0.083] | 1338.017 | 7.101 | 0.001 | 1.49E-03 |
| CASP-8 | -0.384 | -3.471 | 5.39E-04 | [-0.601, -0.167] | -0.378 | -3.266 | 1.12E-03 | [-0.605, -0.151] | 3015.489 | 6.850 | 0.001 | 1.59E-03 |
| ADA | -0.236 | -3.551 | 3.99E-04 | [-0.367, -0.106] | -0.183 | -2.642 | 8.36E-03 | [-0.319, -0.047] | 1847.917 | 6.375 | 0.002 | 2.49E-03 |
| 4E-BP1 | -0.420 | -3.464 | 5.52E-04 | [-0.657, -0.182] | -0.364 | -2.987 | 3.00E-03 | [-0.602, -0.125] | 3220.209 | 6.385 | 0.002 | 2.50E-03 |
| CXCL9 | -0.282 | -2.814 | 4.98E-03 | [-0.479, -0.086] | -0.293 | -3.119 | 2.52E-03 | [-0.478, -0.109] | 2788.670 | 5.261 | 0.007 | 8.87E-03 |
| CCL25 | -0.072 | -1.018 | 3.09E-01 | [-0.211, 0.067] | 0.067 | 1.063 | 2.92E-01 | [-0.056, 0.190] | 1998.789 | 3.844 | 0.026 | 3.43E-02 |
| FGF-21 | -0.039 | -0.163 | 8.70E-01 | [-0.513, 0.434] | 0.446 | 1.830 | 6.79E-02 | [-0.032, 0.924] | 4802.446 | 3.462 | 0.032 | 4.16E-02 |
| ST1A1 | -0.385 | -2.545 | 1.10E-02 | [-0.681, -0.088] | -0.251 | -1.584 | 1.14E-01 | [-0.561, 0.060] | 3732.288 | 3.250 | 0.039 | 5.00E-02 |
| IL-18R1 | 0.021 | 0.369 | 7.12E-01 | [-0.092, 0.135] | 0.111 | 1.910 | 5.70E-02 | [-0.003, 0.226] | 1529.938 | 2.599 | 0.075 | 9.45E-02 |
| Flt3L | 0.083 | 1.633 | 1.03E-01 | [-0.017, 0.183] | 0.111 | 2.265 | 2.48E-02 | [0.015, 0.208] | 1232.909 | 2.572 | 0.079 | 9.67E-02 |
| CST5 | -0.130 | -1.940 | 5.26E-02 | [-0.260, 0.001] | -0.047 | -0.719 | 4.73E-01 | [-0.175, 0.081] | 1854.042 | 2.229 | 0.110 | 1.33E-01 |
| CXCL1 | -0.084 | -0.689 | 4.91E-01 | [-0.321, 0.154] | -0.228 | -1.795 | 7.29E-02 | [-0.477, 0.021] | 3225.681 | 1.838 | 0.160 | 1.90E-01 |
| CXCL11 | -0.208 | -1.704 | 8.86E-02 | [-0.447, 0.031] | -0.189 | -1.490 | 1.36E-01 | [-0.438, 0.060] | 3236.246 | 1.566 | 0.209 | 2.45E-01 |
| CCL3 | 0.077 | 0.866 | 3.86E-01 | [-0.097, 0.251] | 0.145 | 1.613 | 1.07E-01 | [-0.031, 0.322] | 2506.880 | 1.345 | 0.261 | 3.00E-01 |
| CCL4 | -0.102 | -1.409 | 1.59E-01 | [-0.244, 0.040] | -0.095 | -1.278 | 2.02E-01 | [-0.240, 0.051] | 2040.403 | 1.094 | 0.335 | 3.79E-01 |
| CCL20 | -0.165 | -1.336 | 1.82E-01 | [-0.408, 0.077] | -0.087 | -0.839 | 4.05E-01 | [-0.289, 0.116] | 3278.035 | 0.921 | 0.406 | 4.40E-01 |
| IL10 | 0.019 | 0.229 | 8.19E-01 | [-0.145, 0.184] | -0.064 | -0.797 | 4.28E-01 | [-0.220, 0.093] | 2380.127 | 0.903 | 0.408 | 4.40E-01 |
| MMP-1 | 0.037 | 0.268 | 7.89E-01 | [-0.233, 0.307] | 0.167 | 1.161 | 2.46E-01 | [-0.115, 0.449] | 3519.963 | 0.903 | 0.406 | 4.40E-01 |
| FGF-19 | 0.073 | 0.565 | 5.72E-01 | [-0.181, 0.327] | -0.043 | -0.356 | 7.22E-01 | [-0.283, 0.196] | 3378.565 | 0.703 | 0.497 | 5.23E-01 |
| OSM | 0.136 | 1.164 | 2.45E-01 | [-0.093, 0.366] | 0.084 | 0.695 | 4.88E-01 | [-0.153, 0.320] | 3141.095 | 0.683 | 0.505 | 5.23E-01 |
| TNF | -0.063 | -0.958 | 3.38E-01 | [-0.191, 0.066] | -0.011 | -0.162 | 8.71E-01 | [-0.138, 0.117] | 1809.609 | 0.679 | 0.508 | 5.23E-01 |
| IL6 | -0.050 | -0.519 | 6.04E-01 | [-0.241, 0.140] | -0.072 | -0.721 | 4.71E-01 | [-0.268, 0.124] | 2713.506 | 0.261 | 0.770 | 7.82E-01 |
| TGF-α | 0.006 | 0.120 | 9.04E-01 | [-0.092, 0.104] | -0.002 | -0.035 | 9.72E-01 | [-0.104, 0.101] | 1194.707 | 0.019 | 0.981 | 9.81E-01 |

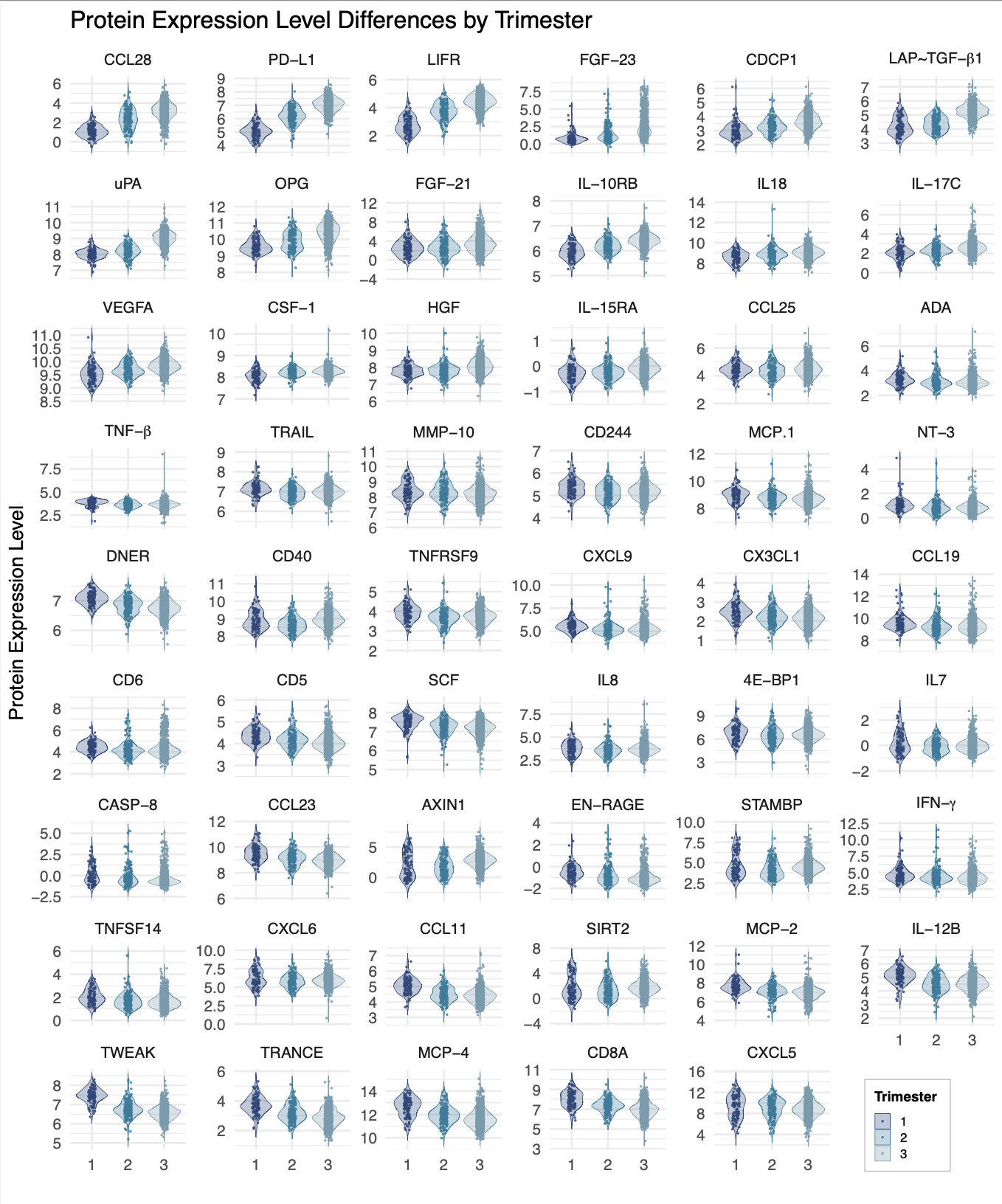

**Supplementary Figure 3. Trimester-related differences in protein expression levels, sorted by significance.** Univariate mixed-effects regression models detected 53 inflammatory proteins that showed significant differences across trimesters at 5% FDR. Models included maternal age at blood draw, gravidity, and infant sex as covariates, with site modeled as a random intercept to account for site-level heterogeneity. Protein expression levels are displayed by trimester (first, second, and third), with violin plots illustrating the distribution of values and overlaid points representing individual samples. Violin plots illustrate clear shifts in the distribution of protein expression levels across the first, second, and third trimesters.

Supplementary Table 4. Main effect of maternal age on expression level for each protein. Abbreviations: AIC, Akaike information criterion; CI, confidence interval; FDR, false discovery rate.

| **Protein** | **B estimate** | **t statistic** | **95% CI** | **AIC** | **p (uncorrected)** | **p (FDR-adjusted)** |
| --- | --- | --- | --- | --- | --- | --- |
| Flt3L | 0.012 | 4.653 | [0.007, 0.017] | 1232.280 | 0.000 | <0.001 |
| FGF-23 | -0.045 | -4.494 | [-0.065, -0.025] | 4327.788 | 0.000 | <0.001 |
| IL-10RB | 0.005 | 3.063 | [0.002, 0.009] | 281.164 | 0.002 | 0.039 |
| PD-L1 | 0.010 | 3.080 | [0.004, 0.017] | 1752.450 | 0.002 | 0.039 |
| CCL20 | -0.017 | -2.882 | [-0.029, -0.006] | 3281.326 | 0.004 | 0.050 |
| FGF-21 | 0.035 | 2.810 | [0.011, 0.059] | 4795.426 | 0.005 | 0.050 |
| MCP-4 | 0.014 | 2.827 | [0.004, 0.024] | 2689.662 | 0.005 | 0.050 |
| FGF-19 | 0.018 | 2.729 | [0.005, 0.031] | 3380.169 | 0.007 | 0.057 |
| LIF-R | 0.009 | 2.600 | [0.002, 0.015] | 1808.548 | 0.010 | 0.071 |
| MCP-1 | 0.008 | 2.579 | [0.002, 0.014] | 1725.321 | 0.010 | 0.071 |
| CD40 | 0.007 | 2.369 | [0.001, 0.013] | 1475.780 | 0.018 | 0.083 |
| CDCP1 | 0.009 | 2.427 | [0.002, 0.015] | 2017.990 | 0.016 | 0.083 |
| CXCL9 | 0.012 | 2.433 | [0.002, 0.022] | 2796.562 | 0.015 | 0.083 |
| HGF | 0.007 | 2.397 | [0.001, 0.012] | 1357.235 | 0.017 | 0.083 |
| SCF | 0.006 | 2.399 | [0.001, 0.011] | 1239.109 | 0.017 | 0.083 |
| VEGFA | 0.004 | 2.296 | [0.001, 0.008] | 286.763 | 0.022 | 0.094 |
| CCL11 | 0.007 | 2.214 | [0.001, 0.013] | 1652.152 | 0.027 | 0.110 |
| CCL25 | 0.008 | 2.180 | [0.001, 0.014] | 1997.570 | 0.030 | 0.114 |
| TRAIL | -0.004 | -2.156 | [-0.008, 0.000] | 637.669 | 0.031 | 0.114 |
| MMP-1 | 0.015 | 2.043 | [0.001, 0.029] | 3522.076 | 0.041 | 0.142 |
| EN-RAGE | -0.009 | -1.932 | [-0.018, 0.000] | 2433.329 | 0.054 | 0.176 |
| CST5 | 0.007 | 1.913 | [0.000, 0.013] | 1858.724 | 0.056 | 0.176 |
| MMP-10 | 0.007 | 1.881 | [0.000, 0.014] | 2127.907 | 0.060 | 0.177 |
| TNFSF14 | 0.007 | 1.871 | [0.000, 0.015] | 2122.743 | 0.062 | 0.177 |
| CD5 | -0.004 | -1.779 | [-0.009, 0.000] | 1000.560 | 0.076 | 0.201 |
| uPA | 0.004 | 1.784 | [0.000, 0.009] | 947.791 | 0.075 | 0.201 |
| TRANCE | -0.006 | -1.725 | [-0.013, 0.001] | 1963.576 | 0.085 | 0.217 |
| CD8A | -0.008 | -1.612 | [-0.018, 0.002] | 2682.904 | 0.107 | 0.264 |
| CSF-1 | 0.002 | 1.526 | [-0.001, 0.005] | -341.897 | 0.127 | 0.274 |
| CXCL6 | 0.009 | 1.565 | [-0.002, 0.020] | 2957.322 | 0.118 | 0.274 |
| DNER | 0.003 | 1.554 | [-0.001, 0.006] | 318.439 | 0.121 | 0.274 |
| IL-18R1 | -0.005 | -1.535 | [-0.010, 0.001] | 1525.699 | 0.125 | 0.274 |
| CASP-8 | 0.009 | 1.457 | [-0.003, 0.020] | 3019.536 | 0.145 | 0.295 |
| IL-17C | -0.007 | -1.469 | [-0.015, 0.002] | 2398.564 | 0.142 | 0.295 |
| IL7 | 0.005 | 1.424 | [-0.002, 0.013] | 1972.134 | 0.155 | 0.298 |
| MCP-2 | 0.007 | 1.422 | [-0.003, 0.017] | 2650.408 | 0.155 | 0.298 |
| IL-15RA | 0.003 | 1.406 | [-0.001, 0.007] | 535.796 | 0.160 | 0.298 |
| OSM | 0.009 | 1.390 | [-0.004, 0.021] | 3143.508 | 0.165 | 0.299 |
| IL8 | 0.006 | 1.288 | [-0.003, 0.015] | 2473.203 | 0.198 | 0.350 |
| AXIN1 | 0.010 | 1.219 | [-0.006, 0.025] | 3720.330 | 0.223 | 0.385 |
| IL6 | -0.006 | -1.169 | [-0.016, 0.004] | 2713.414 | 0.243 | 0.399 |
| NT-3 | -0.004 | -1.178 | [-0.010, 0.003] | 1755.254 | 0.240 | 0.399 |
| CCL23 | 0.004 | 1.120 | [-0.003, 0.010] | 1755.134 | 0.263 | 0.421 |
| CCL3 | 0.005 | 1.093 | [-0.004, 0.014] | 2506.297 | 0.275 | 0.421 |
| CCL4 | 0.004 | 1.101 | [-0.003, 0.012] | 2042.125 | 0.271 | 0.421 |
| CD244 | 0.002 | 0.959 | [-0.002, 0.007] | 981.910 | 0.338 | 0.486 |
| CX3CL1 | 0.003 | 0.962 | [-0.003, 0.008] | 1427.123 | 0.336 | 0.486 |
| IL10 | -0.004 | -0.965 | [-0.012, 0.004] | 2381.891 | 0.336 | 0.486 |
| CCL28 | -0.005 | -0.870 | [-0.016, 0.006] | 3076.633 | 0.385 | 0.532 |
| IL18 | -0.003 | -0.880 | [-0.011, 0.004] | 2094.983 | 0.379 | 0.532 |
| TNFB | -0.002 | -0.851 | [-0.008, 0.003] | 1345.834 | 0.395 | 0.535 |
| TWEAK | 0.002 | 0.748 | [-0.003, 0.007] | 1024.157 | 0.455 | 0.604 |
| TNFRSF9 | -0.002 | -0.727 | [-0.007, 0.003] | 1165.991 | 0.467 | 0.608 |
| CXCL11 | -0.003 | -0.482 | [-0.016, 0.010] | 3240.201 | 0.630 | 0.733 |
| IL-12B | 0.002 | 0.535 | [-0.006, 0.010] | 2141.480 | 0.593 | 0.733 |
| LAP-TGF-β-1 | 0.002 | 0.489 | [-0.005, 0.008] | 1717.636 | 0.625 | 0.733 |
| ST1A1 | 0.004 | 0.472 | [-0.012, 0.019] | 3732.799 | 0.637 | 0.733 |
| STAMBP | 0.004 | 0.557 | [-0.009, 0.016] | 3277.915 | 0.577 | 0.733 |
| TGF-α | -0.001 | -0.483 | [-0.006, 0.004] | 1193.492 | 0.629 | 0.733 |
| TNF | 0.002 | 0.530 | [-0.005, 0.008] | 1811.004 | 0.596 | 0.733 |
| SIRT2 | 0.003 | 0.369 | [-0.013, 0.019] | 3835.821 | 0.712 | 0.806 |
| ADA | 0.001 | 0.251 | [-0.006, 0.008] | 1856.512 | 0.802 | 0.865 |
| CD6 | -0.001 | -0.225 | [-0.009, 0.007] | 2311.437 | 0.822 | 0.865 |
| CXCL5 | 0.002 | 0.219 | [-0.018, 0.022] | 4274.840 | 0.827 | 0.865 |
| OPG | 0.001 | 0.279 | [-0.006, 0.008] | 2021.292 | 0.781 | 0.865 |
| 4E-BP1 | -0.002 | -0.244 | [-0.014, 0.011] | 3226.610 | 0.807 | 0.865 |
| IFN-γ | 0.001 | 0.183 | [-0.012, 0.015] | 3521.805 | 0.855 | 0.881 |
| CXCL1 | 0.001 | 0.158 | [-0.012, 0.014] | 3222.126 | 0.874 | 0.887 |
| CCL19 | 0.000 | 0.026 | [-0.009, 0.009] | 2600.078 | 0.979 | 0.979 |

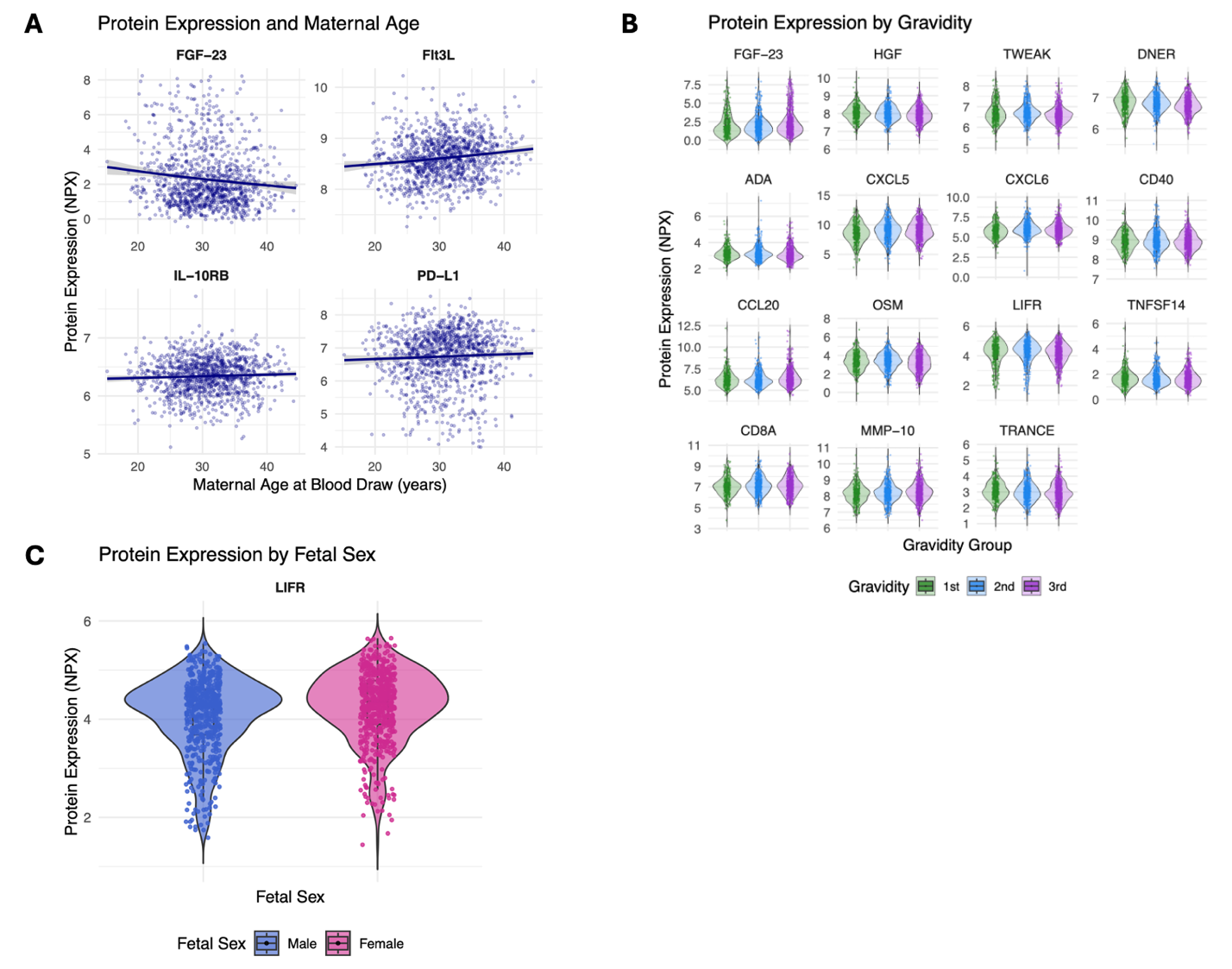

**Supplementary Figure 4. Associations of fetal sex, gravidity, and maternal age with overall protein expression levels.** Scatterplots (bottom left) show linear associations between maternal age at blood draw and protein expression (A). Shaded bands represent 95% confidence intervals. Violin plots display protein expression (Normalized Protein eXpression, NPX) stratified by gravidity group (B) and fetal sex (C). Linear mixed-effects models tested main effects of fetal sex, gravidity, and maternal age on overall protein levels, adjusting for the other covariates as appropriate and including a random intercept for ECHO site. Only proteins with significant associations (p<0.05) are shown. Full model results are provided below. Four proteins showed significant associations with maternal age at blood draw: FGF-23, Flt3L, IL-10RB, and PD-L1. FGF-23 expression decreased with maternal age, while Flt3L, IL-10RB, and PD-L1 expression increased with maternal age (A). We also identified 15 proteins that differed by gravidity status: FGF-23, HGF, TWEAK, DNER, ADA, CXCL5, CXCL6, CD40, CCL20, OSM, LIFR, TNFSF14, CD8A, MMP-10, and TRANCE (B). In contrast, we identified one protein that differed by fetal sex: LIFR expression was higher in pregnancies with female fetuses (C). See Supplementary Tables 4–6 for full results.

Supplementary Table 5. Main effect of gravidity **on expression level for each protein.** Abbreviations: AIC, Akaike information criterion; CI, confidence interval; FDR, false discovery rate.

| **Protein** | **G1–G2**  **β estimate** | **G1–G2**  **t statistic** | **G1–G2 p** | **G1–G2 95% CI** | **G1–G3**  **B estimate** | **G1–G3**  **t statistic** | **G1–G3 p** | **G1–G3 95% CI** | **AIC** | **Omnibus F** | **Omnibus p (uncorrected)** | **Omnibus p (FDR-adjusted)** |
| --- | --- | --- | --- | --- | --- | --- | --- | --- | --- | --- | --- | --- |
| HGF | -0.066 | -1.799 | 0.072 | [-0.138, 0.006] | -0.148 | -4.131 | 0.000 | [-0.219, -0.078] | 1357.235 | 9.107 | <0.001 | 0.003 |
| FGF-23 | -0.021 | -0.156 | 0.876 | [-0.283, 0.241] | 0.416 | 3.219 | 0.001 | [0.163, 0.669] | 4327.788 | 9.285 | <0.001 | 0.003 |
| TWEAK | 0.021 | 0.655 | 0.513 | [-0.042, 0.083] | -0.089 | -2.858 | 0.004 | [-0.150, -0.028] | 1024.157 | 9.200 | <0.001 | 0.003 |
| DNER | -0.020 | -0.874 | 0.382 | [-0.066, 0.025] | -0.080 | -3.526 | 0.000 | [-0.124, -0.035] | 318.439 | 7.796 | <0.001 | 0.008 |
| ADA | 0.018 | 0.391 | 0.696 | [-0.072, 0.108] | -0.120 | -2.670 | 0.008 | [-0.209, -0.032] | 1856.512 | 7.198 | <0.001 | 0.011 |
| CD40 | -0.007 | -0.177 | 0.860 | [-0.083, 0.069] | -0.106 | -2.775 | 0.006 | [-0.181, -0.031] | 1475.780 | 5.942 | 0.003 | 0.026 |
| CCL20 | 0.011 | 0.142 | 0.887 | [-0.145, 0.168] | 0.206 | 2.683 | 0.007 | [0.055, 0.356] | 3281.326 | 5.725 | 0.003 | 0.026 |
| CXCL6 | 0.186 | 2.505 | 0.012 | [0.040, 0.331] | -0.012 | -0.168 | 0.867 | [-0.155, 0.131] | 2957.322 | 5.718 | 0.003 | 0.026 |
| CXCL5 | 0.339 | 2.577 | 0.010 | [0.081, 0.597] | -0.023 | -0.181 | 0.856 | [-0.277, 0.230] | 4274.840 | 6.076 | 0.002 | 0.026 |
| OSM | -0.063 | -0.781 | 0.435 | [-0.220, 0.095] | -0.237 | -3.004 | 0.003 | [-0.391, -0.082] | 3143.508 | 5.573 | 0.004 | 0.027 |
| LIF-R | -0.022 | -0.489 | 0.625 | [-0.110, 0.066] | -0.122 | -2.813 | 0.005 | [-0.207, -0.037] | 1808.548 | 5.378 | 0.005 | 0.030 |
| TNFSF14 | 0.014 | 0.273 | 0.785 | [-0.087, 0.115] | -0.114 | -2.249 | 0.025 | [-0.213, -0.015] | 2122.743 | 4.942 | 0.007 | 0.042 |
| MMP-10 | 0.053 | 1.101 | 0.271 | [-0.042, 0.148] | 0.136 | 2.926 | 0.004 | [0.045, 0.227] | 2127.907 | 4.793 | 0.008 | 0.043 |
| CD8A | 0.168 | 2.561 | 0.011 | [0.040, 0.297] | 0.191 | 2.958 | 0.003 | [0.064, 0.318] | 2682.904 | 4.767 | 0.009 | 0.043 |
| TRANCE | -0.074 | -1.546 | 0.122 | [-0.167, 0.020] | -0.139 | -3.000 | 0.003 | [-0.230, -0.048] | 1963.576 | 4.610 | 0.010 | 0.047 |
| CX3CL1 | -0.050 | -1.322 | 0.186 | [-0.125, 0.024] | -0.106 | -2.861 | 0.004 | [-0.178, -0.033] | 1427.123 | 4.298 | 0.014 | 0.060 |
| CD244 | 0.062 | 1.990 | 0.047 | [0.001, 0.124] | -0.011 | -0.364 | 0.716 | [-0.072, 0.049] | 981.910 | 4.194 | 0.015 | 0.062 |
| IL7 | 0.092 | 1.913 | 0.056 | [-0.002, 0.187] | -0.021 | -0.440 | 0.660 | [-0.114, 0.072] | 1972.134 | 4.116 | 0.017 | 0.063 |
| TNFRSF9 | 0.011 | 0.333 | 0.739 | [-0.055, 0.078] | -0.063 | -1.901 | 0.058 | [-0.128, 0.002] | 1165.991 | 3.798 | 0.023 | 0.083 |
| CASP-8 | 0.104 | 1.360 | 0.174 | [-0.046, 0.253] | -0.070 | -0.934 | 0.351 | [-0.217, 0.077] | 3019.536 | 3.622 | 0.027 | 0.093 |
| uPA | 0.006 | 0.190 | 0.849 | [-0.055, 0.066] | -0.058 | -1.905 | 0.057 | [-0.117, 0.002] | 947.791 | 3.451 | 0.032 | 0.105 |
| CXCL1 | 0.193 | 2.325 | 0.020 | [0.030, 0.356] | 0.047 | 0.575 | 0.565 | [-0.113, 0.207] | 3222.126 | 3.377 | 0.034 | 0.108 |
| CST5 | -0.062 | -1.369 | 0.171 | [-0.152, 0.027] | -0.109 | -2.464 | 0.014 | [-0.195, -0.022] | 1858.724 | 3.068 | 0.047 | 0.141 |
| Flt3L | 0.068 | 1.965 | 0.050 | [0.000, 0.136] | 0.005 | 0.138 | 0.890 | [-0.061, 0.070] | 1232.280 | 2.976 | 0.052 | 0.148 |
| IL-10RB | -0.002 | -0.098 | 0.922 | [-0.048, 0.043] | -0.042 | -1.841 | 0.066 | [-0.086, 0.003] | 281.164 | 2.653 | 0.071 | 0.188 |
| IL18 | 0.084 | 1.648 | 0.100 | [-0.016, 0.183] | 0.114 | 2.306 | 0.021 | [0.017, 0.211] | 2094.983 | 2.687 | 0.069 | 0.188 |
| OPG | 0.040 | 0.829 | 0.408 | [-0.055, 0.135] | 0.100 | 2.155 | 0.032 | [0.009, 0.192] | 2021.292 | 2.578 | 0.077 | 0.198 |
| MCP-4 | -0.058 | -0.873 | 0.383 | [-0.187, 0.072] | -0.136 | -2.098 | 0.036 | [-0.263, -0.009] | 2689.662 | 2.375 | 0.093 | 0.230 |
| SCF | 0.002 | 0.055 | 0.956 | [-0.063, 0.067] | -0.052 | -1.619 | 0.107 | [-0.114, 0.011] | 1239.109 | 2.331 | 0.100 | 0.238 |
| NT-3 | -0.030 | -0.701 | 0.484 | [-0.115, 0.055] | -0.083 | -2.000 | 0.046 | [-0.165, -0.002] | 1755.254 | 2.279 | 0.103 | 0.238 |
| IFN-γ | -0.156 | -1.670 | 0.095 | [-0.338, 0.027] | -0.012 | -0.137 | 0.891 | [-0.188, 0.163] | 3521.805 | 2.124 | 0.121 | 0.269 |
| PD-L1 | -0.017 | -0.390 | 0.697 | [-0.103, 0.069] | -0.077 | -1.788 | 0.074 | [-0.161, 0.007] | 1752.450 | 2.065 | 0.127 | 0.275 |
| CCL28 | 0.019 | 0.244 | 0.807 | [-0.130, 0.167] | 0.121 | 1.671 | 0.096 | [-0.021, 0.264] | 3076.633 | 1.975 | 0.141 | 0.294 |
| LAP-TGF-β-1 | -0.037 | -0.860 | 0.390 | [-0.122, 0.048] | -0.079 | -1.857 | 0.064 | [-0.162, 0.004] | 1717.636 | 1.808 | 0.164 | 0.334 |
| TGF-α | -0.050 | -1.467 | 0.143 | [-0.118, 0.017] | -0.059 | -1.744 | 0.081 | [-0.125, 0.007] | 1193.492 | 1.630 | 0.196 | 0.387 |
| TRAIL | 0.037 | 1.388 | 0.166 | [-0.015, 0.090] | 0.000 | 0.011 | 0.991 | [-0.051, 0.052] | 637.669 | 1.597 | 0.203 | 0.389 |
| TNF | 0.018 | 0.409 | 0.682 | [-0.070, 0.106] | 0.068 | 1.567 | 0.118 | [-0.017, 0.154] | 1811.004 | 1.516 | 0.220 | 0.410 |
| IL8 | 0.100 | 1.667 | 0.096 | [-0.018, 0.218] | 0.066 | 1.129 | 0.259 | [-0.049, 0.182] | 2473.203 | 1.394 | 0.249 | 0.451 |
| CCL4 | -0.064 | -1.296 | 0.195 | [-0.162, 0.033] | -0.076 | -1.567 | 0.117 | [-0.172, 0.019] | 2042.125 | 1.304 | 0.272 | 0.469 |
| CCL3 | -0.049 | -0.809 | 0.419 | [-0.168, 0.070] | 0.034 | 0.575 | 0.565 | [-0.082, 0.151] | 2506.297 | 1.321 | 0.267 | 0.469 |
| IL6 | 0.003 | 0.049 | 0.961 | [-0.127, 0.134] | 0.080 | 1.234 | 0.218 | [-0.047, 0.208] | 2713.414 | 1.212 | 0.298 | 0.501 |
| STAMBP | 0.068 | 0.795 | 0.427 | [-0.099, 0.235] | -0.039 | -0.462 | 0.644 | [-0.203, 0.126] | 3277.915 | 1.097 | 0.334 | 0.549 |
| SIRT2 | 0.052 | 0.474 | 0.635 | [-0.161, 0.264] | -0.077 | -0.719 | 0.472 | [-0.286, 0.133] | 3835.821 | 0.981 | 0.375 | 0.602 |
| IL-12B | -0.040 | -0.778 | 0.437 | [-0.142, 0.061] | -0.067 | -1.320 | 0.187 | [-0.167, 0.033] | 2141.480 | 0.874 | 0.417 | 0.643 |
| FGF-19 | -0.031 | -0.358 | 0.721 | [-0.203, 0.141] | -0.098 | -1.165 | 0.244 | [-0.264, 0.067] | 3380.169 | 0.806 | 0.447 | 0.643 |
| CCL19 | -0.024 | -0.380 | 0.704 | [-0.148, 0.100] | -0.072 | -1.178 | 0.239 | [-0.192, 0.048] | 2600.078 | 0.809 | 0.446 | 0.643 |
| CSF-1 | 0.018 | 1.037 | 0.300 | [-0.016, 0.053] | 0.001 | 0.037 | 0.970 | [-0.033, 0.034] | -341.897 | 0.861 | 0.423 | 0.643 |
| AXIN1 | 0.054 | 0.525 | 0.600 | [-0.148, 0.256] | -0.059 | -0.577 | 0.564 | [-0.258, 0.140] | 3720.330 | 0.830 | 0.436 | 0.643 |
| CCL25 | -0.051 | -1.103 | 0.272 | [-0.142, 0.040] | -0.044 | -0.996 | 0.320 | [-0.131, 0.043] | 1997.570 | 0.679 | 0.508 | 0.662 |
| MCP-1 | -0.023 | -0.536 | 0.592 | [-0.105, 0.060] | -0.047 | -1.174 | 0.241 | [-0.127, 0.032] | 1725.321 | 0.729 | 0.483 | 0.662 |
| CD5 | 0.037 | 1.176 | 0.240 | [-0.025, 0.099] | 0.031 | 0.990 | 0.323 | [-0.030, 0.092] | 1000.560 | 0.741 | 0.477 | 0.662 |
| IL-18R1 | 0.044 | 1.116 | 0.265 | [-0.033, 0.122] | 0.037 | 0.967 | 0.334 | [-0.038, 0.113] | 1525.699 | 0.678 | 0.508 | 0.662 |
| MMP-1 | 0.093 | 0.986 | 0.324 | [-0.092, 0.279] | 0.012 | 0.128 | 0.898 | [-0.171, 0.195] | 3522.076 | 0.693 | 0.500 | 0.662 |
| IL-15RA | -0.010 | -0.400 | 0.689 | [-0.061, 0.040] | 0.014 | 0.569 | 0.569 | [-0.035, 0.064] | 535.796 | 0.648 | 0.523 | 0.663 |
| CD6 | 0.060 | 1.074 | 0.283 | [-0.049, 0.169] | 0.023 | 0.411 | 0.681 | [-0.085, 0.130] | 2311.437 | 0.638 | 0.529 | 0.663 |
| ST1A1 | 0.105 | 1.016 | 0.310 | [-0.098, 0.309] | 0.036 | 0.354 | 0.723 | [-0.164, 0.236] | 3732.799 | 0.586 | 0.556 | 0.686 |
| FGF-21 | -0.016 | -0.097 | 0.923 | [-0.338, 0.306] | 0.113 | 0.707 | 0.480 | [-0.200, 0.426] | 4795.426 | 0.501 | 0.606 | 0.733 |
| VEGFA | -0.001 | -0.022 | 0.982 | [-0.046, 0.045] | -0.017 | -0.735 | 0.463 | [-0.061, 0.028] | 286.763 | 0.435 | 0.648 | 0.733 |
| TNFB | 0.009 | 0.246 | 0.806 | [-0.062, 0.080] | -0.019 | -0.545 | 0.586 | [-0.088, 0.050] | 1345.834 | 0.446 | 0.641 | 0.733 |
| CXCL9 | 0.018 | 0.266 | 0.791 | [-0.115, 0.152] | 0.058 | 0.887 | 0.376 | [-0.070, 0.187] | 2796.562 | 0.470 | 0.625 | 0.733 |
| CXCL11 | 0.077 | 0.920 | 0.358 | [-0.087, 0.241] | 0.039 | 0.469 | 0.639 | [-0.123, 0.200] | 3240.201 | 0.434 | 0.648 | 0.733 |
| CCL23 | 0.026 | 0.585 | 0.559 | [-0.060, 0.112] | -0.004 | -0.098 | 0.922 | [-0.089, 0.080] | 1755.134 | 0.355 | 0.701 | 0.780 |
| MCP-2 | -0.050 | -0.776 | 0.438 | [-0.177, 0.077] | -0.044 | -0.699 | 0.485 | [-0.168, 0.080] | 2650.408 | 0.336 | 0.714 | 0.782 |
| IL-17C | -0.038 | -0.658 | 0.510 | [-0.152, 0.076] | -0.036 | -0.635 | 0.525 | [-0.148, 0.075] | 2398.564 | 0.256 | 0.774 | 0.826 |
| CCL11 | -0.030 | -0.706 | 0.480 | [-0.111, 0.052] | -0.017 | -0.407 | 0.684 | [-0.096, 0.063] | 1652.152 | 0.251 | 0.778 | 0.826 |
| 4E-BP1 | 0.025 | 0.296 | 0.768 | [-0.139, 0.188] | -0.023 | -0.287 | 0.774 | [-0.183, 0.136] | 3226.610 | 0.233 | 0.792 | 0.828 |
| CDCP1 | 0.006 | 0.134 | 0.894 | [-0.086, 0.099] | -0.016 | -0.363 | 0.717 | [-0.105, 0.072] | 2017.990 | 0.180 | 0.835 | 0.860 |
| IL10 | -0.023 | -0.409 | 0.683 | [-0.132, 0.086] | -0.028 | -0.528 | 0.598 | [-0.132, 0.076] | 2381.891 | 0.143 | 0.866 | 0.879 |
| EN-RAGE | 0.018 | 0.310 | 0.756 | [-0.097, 0.134] | 0.000 | 0.004 | 0.997 | [-0.113, 0.114] | 2433.329 | 0.079 | 0.924 | 0.924 |

Supplementary Table 6. Main effect of fetal sex on expression level for each protein. Abbreviations: AIC, Akaike information criterion; CI, confidence interval; FDR, false discovery rate.

| **Protein** | **B estimate** | **t statistic** | **95% CI** | **AIC** | **p (uncorrected)** | **p (FDR-adjusted)** |
| --- | --- | --- | --- | --- | --- | --- |
| LIF-R | 0.135 | 4.415 | [0.075, 0.194] | 1808.548 | 0.000 | 0.001 |
| CD244 | -0.061 | -2.884 | [-0.103, -0.020] | 981.910 | 0.004 | 0.081 |
| IL-10RB | -0.047 | -3.030 | [-0.078, -0.017] | 281.164 | 0.002 | 0.081 |
| IL10 | -0.111 | -2.834 | [-0.188, -0.034] | 2381.891 | 0.005 | 0.081 |
| IL-18R1 | -0.066 | -2.444 | [-0.119, -0.013] | 1525.699 | 0.015 | 0.191 |
| TNFRSF9 | -0.055 | -2.399 | [-0.100, -0.010] | 1165.991 | 0.017 | 0.191 |
| PD-L1 | 0.069 | 2.336 | [0.011, 0.128] | 1752.450 | 0.020 | 0.194 |
| CD40 | -0.060 | -2.284 | [-0.111, -0.009] | 1475.780 | 0.023 | 0.194 |
| IL-15RA | -0.038 | -2.158 | [-0.072, -0.003] | 535.796 | 0.031 | 0.239 |
| CD6 | -0.070 | -1.850 | [-0.144, 0.004] | 2311.437 | 0.065 | 0.446 |
| CCL25 | -0.055 | -1.664 | [-0.120, 0.010] | 1997.570 | 0.096 | 0.514 |
| CXCL9 | -0.078 | -1.662 | [-0.170, 0.014] | 2796.562 | 0.097 | 0.514 |
| TRANCE | -0.056 | -1.728 | [-0.120, 0.008] | 1963.576 | 0.084 | 0.514 |
| CD5 | -0.033 | -1.544 | [-0.075, 0.009] | 1000.560 | 0.123 | 0.529 |
| FGF-23 | 0.142 | 1.549 | [-0.038, 0.321] | 4327.788 | 0.122 | 0.529 |
| IFN-γ | -0.102 | -1.585 | [-0.228, 0.024] | 3521.805 | 0.113 | 0.529 |
| LAP-TGF-β-1 | -0.043 | -1.476 | [-0.100, 0.014] | 1717.636 | 0.140 | 0.569 |
| CASP-8 | 0.072 | 1.404 | [-0.029, 0.173] | 3019.536 | 0.161 | 0.589 |
| MMP-10 | -0.049 | -1.399 | [-0.118, 0.020] | 2127.907 | 0.162 | 0.589 |
| CXCL5 | 0.116 | 1.301 | [-0.058, 0.290] | 4274.840 | 0.193 | 0.635 |
| IL18 | -0.045 | -1.311 | [-0.113, 0.022] | 2094.983 | 0.190 | 0.635 |
| CXCL1 | 0.064 | 1.139 | [-0.046, 0.174] | 3222.126 | 0.255 | 0.705 |
| IL-12B | -0.042 | -1.183 | [-0.111, 0.027] | 2141.480 | 0.237 | 0.705 |
| IL-17C | 0.045 | 1.138 | [-0.032, 0.122] | 2398.564 | 0.255 | 0.705 |
| uPA | -0.025 | -1.210 | [-0.066, 0.016] | 947.791 | 0.227 | 0.705 |
| HGF | -0.028 | -1.100 | [-0.077, 0.022] | 1357.235 | 0.271 | 0.719 |
| OSM | 0.059 | 1.078 | [-0.048, 0.165] | 3143.508 | 0.281 | 0.719 |
| MCP-2 | -0.045 | -1.035 | [-0.132, 0.041] | 2650.408 | 0.301 | 0.742 |
| AXIN1 | -0.069 | -0.989 | [-0.205, 0.068] | 3720.330 | 0.323 | 0.766 |
| FGF-19 | 0.059 | 0.969 | [-0.060, 0.177] | 3380.169 | 0.333 | 0.766 |
| CCL23 | -0.028 | -0.932 | [-0.086, 0.031] | 1755.134 | 0.351 | 0.782 |
| CCL19 | -0.032 | -0.747 | [-0.117, 0.052] | 2600.078 | 0.455 | 0.873 |
| CD8A | 0.036 | 0.820 | [-0.051, 0.124] | 2682.904 | 0.412 | 0.873 |
| CXCL11 | -0.043 | -0.766 | [-0.154, 0.068] | 3240.201 | 0.444 | 0.873 |
| EN-RAGE | 0.031 | 0.778 | [-0.047, 0.109] | 2433.329 | 0.437 | 0.873 |
| TWEAK | -0.017 | -0.768 | [-0.059, 0.026] | 1024.157 | 0.443 | 0.873 |
| DNER | -0.011 | -0.685 | [-0.042, 0.020] | 318.439 | 0.493 | 0.901 |
| FGF-21 | -0.074 | -0.664 | [-0.294, 0.145] | 4795.426 | 0.507 | 0.901 |
| IL8 | 0.027 | 0.660 | [-0.053, 0.106] | 2473.203 | 0.509 | 0.901 |
| CDCP1 | -0.020 | -0.591 | [-0.085, 0.046] | 2017.990 | 0.555 | 0.934 |
| ST1A1 | 0.043 | 0.608 | [-0.095, 0.180] | 3732.799 | 0.543 | 0.934 |
| IL6 | 0.025 | 0.546 | [-0.064, 0.113] | 2713.414 | 0.585 | 0.938 |
| NT-3 | -0.015 | -0.508 | [-0.074, 0.043] | 1755.254 | 0.611 | 0.938 |
| TNF | -0.015 | -0.508 | [-0.075, 0.044] | 1811.004 | 0.612 | 0.938 |
| TNFB | -0.014 | -0.543 | [-0.062, 0.035] | 1345.834 | 0.587 | 0.938 |
| TRAIL | -0.009 | -0.486 | [-0.045, 0.027] | 637.669 | 0.627 | 0.941 |
| ADA | 0.012 | 0.380 | [-0.049, 0.073] | 1856.512 | 0.704 | 0.948 |
| CCL20 | -0.013 | -0.218 | [-0.126, 0.101] | 3281.326 | 0.827 | 0.948 |
| CCL28 | -0.010 | -0.187 | [-0.114, 0.094] | 3076.633 | 0.852 | 0.948 |
| CCL4 | 0.006 | 0.169 | [-0.060, 0.072] | 2042.125 | 0.866 | 0.948 |
| CSF-1 | 0.005 | 0.438 | [-0.018, 0.029] | -341.897 | 0.662 | 0.948 |
| CST5 | -0.010 | -0.335 | [-0.072, 0.051] | 1858.724 | 0.737 | 0.948 |
| CX3CL1 | -0.008 | -0.319 | [-0.059, 0.042] | 1427.123 | 0.749 | 0.948 |
| CXCL6 | -0.010 | -0.204 | [-0.108, 0.088] | 2957.322 | 0.839 | 0.948 |
| Flt3L | 0.008 | 0.354 | [-0.038, 0.055] | 1232.280 | 0.723 | 0.948 |
| IL7 | -0.015 | -0.458 | [-0.079, 0.049] | 1972.134 | 0.647 | 0.948 |
| MCP-1 | 0.006 | 0.219 | [-0.051, 0.064] | 1725.321 | 0.826 | 0.948 |
| MMP-1 | 0.018 | 0.275 | [-0.108, 0.143] | 3522.076 | 0.783 | 0.948 |
| SCF | -0.004 | -0.179 | [-0.051, 0.042] | 1239.109 | 0.858 | 0.948 |
| SIRT2 | -0.016 | -0.212 | [-0.159, 0.128] | 3835.821 | 0.832 | 0.948 |
| STAMBP | -0.018 | -0.310 | [-0.131, 0.095] | 3277.915 | 0.756 | 0.948 |
| TGF-α | 0.005 | 0.234 | [-0.040, 0.051] | 1193.492 | 0.815 | 0.948 |
| VEGFA | -0.007 | -0.417 | [-0.037, 0.024] | 286.763 | 0.677 | 0.948 |
| CCL11 | 0.003 | 0.103 | [-0.053, 0.059] | 1652.152 | 0.918 | 0.951 |
| CCL3 | 0.005 | 0.111 | [-0.076, 0.085] | 2506.297 | 0.912 | 0.951 |
| MCP-4 | 0.005 | 0.110 | [-0.082, 0.092] | 2689.662 | 0.912 | 0.951 |
| OPG | -0.003 | -0.097 | [-0.069, 0.062] | 2021.292 | 0.923 | 0.951 |
| 4E-BP1 | 0.003 | 0.053 | [-0.108, 0.114] | 3226.610 | 0.958 | 0.972 |
| TNFSF14 | -0.001 | -0.031 | [-0.069, 0.067] | 2122.743 | 0.975 | 0.975 |

Supplementary Table 7. Linear models for continuous gestational age. Abbreviations: AIC, Akaike information criterion; CI, confidence interval; FDR, false discovery rate.

| **Protein** | **B estimate** | **t statistic** | **95% CI** | **p (uncorrected)** | **p (FDR-adjusted)** | **Model fit** |
| --- | --- | --- | --- | --- | --- | --- |
| CCL28 | 0.074 | 22.948 | [0.067, 0.080] | 4.12E-09 | 1.14E-08 | AIC=3042.2; R2m=0.324; R2c=0.324 |
| PD-L1 | 0.072 | 27.334 | [0.067, 0.077] | 5.66E-79 | 3.90E-77 | AIC=1714.1; R2m=0.581; R2c=0.602 |
| FGF-23 | 0.065 | 9.829 | [0.052, 0.079] | 4.97E-06 | 1.04E-05 | AIC=4301.3; R2m=0.143; R2c=0.148 |
| LIF-R | 0.065 | 25.677 | [0.060, 0.070] | 5.07E-43 | 1.17E-41 | AIC=1769.6; R2m=0.526; R2c=0.536 |
| FGF-21 | 0.043 | 4.702 | [0.025, 0.061] | 1.44E-05 | 2.83E-05 | AIC=4772.1; R2m=0.043; R2c=0.062 |
| uPA | 0.039 | 20.028 | [0.035, 0.043] | 7.54E-74 | 2.60E-72 | AIC=906.6; R2m=0.419; R2c=0.513 |
| LAP-TGF-β1 | 0.038 | 14.079 | [0.033, 0.044] | 8.76E-41 | 1.51E-39 | AIC=1680.5; R2m=0.261; R2c=0.385 |
| OPG | 0.036 | 17.408 | [0.032, 0.040] | 8.74E-07 | 2.15E-06 | AIC=1981.7; R2m=0.228; R2c=0.229 |
| CDCP1 | 0.029 | 14.236 | [0.025, 0.033] | 9.75E-12 | 3.96E-11 | AIC=1977.7; R2m=0.155; R2c=0.155 |
| IL-17C | 0.020 | 5.529 | [0.013, 0.027] | 5.56E-08 | 1.42E-07 | AIC=2364.1; R2m=0.058; R2c=0.129 |
| IL-10RB | 0.017 | 12.111 | [0.014, 0.020] | 2.29E-28 | 2.25E-27 | AIC=235.4; R2m=0.222; R2c=0.271 |
| HGF | 0.014 | 6.424 | [0.010, 0.018] | 1.81E-09 | 5.67E-09 | AIC=1316.4; R2m=0.083; R2c=0.114 |
| IL18 | 0.013 | 4.536 | [0.008, 0.019] | 1.26E-05 | 2.56E-05 | AIC=2058.0; R2m=0.043; R2c=0.073 |
| VEGFA | 0.011 | 7.298 | [0.008, 0.013] | 8.12E-13 | 4.00E-12 | AIC=241.7; R2m=0.093; R2c=0.199 |
| IL-15RA | 0.009 | 5.965 | [0.006, 0.012] | 1.23E-08 | 3.26E-08 | AIC=490.8; R2m=0.065; R2c=0.101 |
| IL-18R1 | 0.008 | 3.589 | [0.004, 0.012] | 6.44E-04 | 1.03E-03 | AIC=1485.4; R2m=0.029; R2c=0.050 |
| CCL3 | 0.008 | 2.136 | [0.001, 0.015] | 3.43E-02 | 4.30E-02 | AIC=2471.7; R2m=0.012; R2c=0.051 |
| CCL25 | 0.007 | 3.757 | [0.004, 0.011] | 1.81E-04 | 3.04E-04 | AIC=1956.8; R2m=0.018; R2c=0.018 |
| CSF-1 | 0.007 | 6.640 | [0.005, 0.009] | 9.30E-11 | 3.38E-10 | AIC=-390.7; R2m=0.080; R2c=0.150 |
| MMP-1 | 0.007 | 1.194 | [-0.005, 0.019] | 2.33E-01 | 2.64E-01 | AIC=3494.3; R2m=0.008; R2c=0.175 |
| IL6 | 0.005 | 1.350 | [-0.002, 0.013] | 1.78E-01 | 2.05E-01 | AIC=2680.3; R2m=0.008; R2c=0.066 |
| Flt3L | 0.005 | 2.584 | [0.001, 0.008] | 1.26E-02 | 1.67E-02 | AIC=1189.7; R2m=0.036; R2c=0.047 |
| OSM | 0.003 | 0.539 | [-0.007, 0.012] | 5.90E-01 | 6.17E-01 | AIC=3113.0; R2m=0.013; R2c=0.092 |
| IL10 | 0.003 | 1.129 | [-0.002, 0.007] | 2.59E-01 | 2.88E-01 | AIC=2343.3; R2m=0.009; R2c=0.009 |
| TGF-α | 0.002 | 0.944 | [-0.002, 0.006] | 3.45E-01 | 3.78E-01 | AIC=1153.9; R2m=0.005; R2c=0.221 |
| TNF | 0.001 | 0.442 | [-0.004, 0.006] | 6.59E-01 | 6.79E-01 | AIC=1772.3; R2m=0.004; R2c=0.030 |
| CST5 | 0.000 | -0.087 | [-0.005, 0.005] | 9.32E-01 | 9.32E-01 | AIC=1819.5; R2m=0.007; R2c=0.019 |
| CCL20 | -0.001 | -0.401 | [-0.008, 0.005] | 6.88E-01 | 6.98E-01 | AIC=3247.2; R2m=0.015; R2c=0.015 |
| CCL4 | -0.003 | -0.877 | [-0.009, 0.003] | 3.81E-01 | 4.11E-01 | AIC=2005.5; R2m=0.005; R2c=0.062 |
| CXCL11 | -0.004 | -0.688 | [-0.014, 0.007] | 4.91E-01 | 5.22E-01 | AIC=3210.8; R2m=0.002; R2c=0.145 |
| TNFB | -0.005 | -2.577 | [-0.009, -0.001] | 1.34E-02 | 1.75E-02 | AIC=1303.8; R2m=0.011; R2c=0.021 |
| NT-3 | -0.006 | -2.637 | [-0.010, -0.001] | 1.41E-02 | 1.80E-02 | AIC=1714.9; R2m=0.016; R2c=0.022 |
| TNFRSF9 | -0.006 | -2.872 | [-0.010, -0.002] | 4.55E-03 | 6.54E-03 | AIC=1124.4; R2m=0.029; R2c=0.069 |
| CXCL9 | -0.006 | -1.841 | [-0.012, 0.000] | 9.30E-02 | 1.11E-01 | AIC=2761.5; R2m=0.014; R2c=0.017 |
| ADA | -0.006 | -2.049 | [-0.012, -0.000] | 4.07E-02 | 5.02E-02 | AIC=1819.9; R2m=0.021; R2c=0.165 |
| TRAIL | -0.006 | -3.761 | [-0.009, -0.003] | 2.10E-04 | 3.45E-04 | AIC=593.5; R2m=0.033; R2c=0.082 |
| MCP-1 | -0.006 | -3.268 | [-0.010, -0.003] | 3.75E-03 | 5.50E-03 | AIC=1684.1; R2m=0.018; R2c=0.020 |
| FGF-19 | -0.006 | -1.497 | [-0.015, 0.002] | 1.49E-01 | 1.74E-01 | AIC=3348.2; R2m=0.012; R2c=0.017 |
| IL8 | -0.007 | -1.856 | [-0.014, 0.000] | 6.41E-02 | 7.76E-02 | AIC=2439.2; R2m=0.011; R2c=0.091 |
| MMP-10 | -0.009 | -4.150 | [-0.013, -0.005] | 3.57E-05 | 6.49E-05 | AIC=2087.8; R2m=0.029; R2c=0.029 |
| CCL19 | -0.009 | -2.687 | [-0.016, -0.003] | 1.02E-02 | 1.37E-02 | AIC=2565.1; R2m=0.013; R2c=0.029 |
| CD40 | -0.010 | -3.911 | [-0.015, -0.005] | 9.77E-05 | 1.69E-04 | AIC=1438.0; R2m=0.040; R2c=0.305 |
| CX3CL1 | -0.010 | -4.393 | [-0.014, -0.005] | 2.57E-05 | 4.80E-05 | AIC=1386.6; R2m=0.042; R2c=0.070 |
| DNER | -0.011 | -8.620 | [-0.013, -0.008] | 5.38E-12 | 2.32E-11 | AIC=271.4; R2m=0.120; R2c=0.133 |
| SCF | -0.012 | -8.323 | [-0.015, -0.009] | 2.40E-16 | 1.84E-15 | AIC=1194.4; R2m=0.066; R2c=0.066 |
| CD244 | -0.012 | -6.076 | [-0.016, -0.008] | 2.32E-09 | 6.96E-09 | AIC=940.3; R2m=0.079; R2c=0.168 |
| 4E-BP1 | -0.014 | -2.781 | [-0.024, -0.004] | 5.96E-03 | 8.22E-03 | AIC=3195.9; R2m=0.014; R2c=0.059 |
| IL7 | -0.014 | -4.615 | [-0.020, -0.008] | 4.43E-06 | 9.55E-06 | AIC=1936.7; R2m=0.043; R2c=0.262 |
| CD6 | -0.014 | -4.022 | [-0.021, -0.007] | 6.14E-05 | 1.09E-04 | AIC=2279.0; R2m=0.019; R2c=0.531 |
| CD5 | -0.015 | -7.311 | [-0.019, -0.011] | 5.26E-13 | 2.79E-12 | AIC=960.1; R2m=0.085; R2c=0.308 |
| IFN-γ | -0.015 | -3.446 | [-0.024, -0.007] | 2.88E-03 | 4.41E-03 | AIC=3490.5; R2m=0.020; R2c=0.024 |
| CXCL1 | -0.016 | -2.924 | [-0.026, -0.005] | 3.53E-03 | 5.29E-03 | AIC=3193.6; R2m=0.018; R2c=0.318 |
| CASP-8 | -0.016 | -3.291 | [-0.026, -0.006] | 1.03E-03 | 1.62E-03 | AIC=2989.9; R2m=0.024; R2c=0.311 |
| ST1A1 | -0.019 | -2.786 | [-0.032, -0.005] | 5.42E-03 | 7.63E-03 | AIC=3707.3; R2m=0.011; R2c=0.387 |
| CCL11 | -0.019 | -8.239 | [-0.024, -0.015] | 2.50E-11 | 9.57E-11 | AIC=1612.5; R2m=0.106; R2c=0.125 |
| TNFSF14 | -0.021 | -6.340 | [-0.027, -0.014] | 3.52E-10 | 1.16E-09 | AIC=2087.9; R2m=0.076; R2c=0.257 |
| CCL23 | -0.022 | -7.969 | [-0.027, -0.016] | 1.21E-14 | 8.37E-14 | AIC=1717.3; R2m=0.109; R2c=0.181 |
| CXCL6 | -0.022 | -4.675 | [-0.031, -0.013] | 3.37E-06 | 7.51E-06 | AIC=2926.8; R2m=0.048; R2c=0.225 |
| MCP-2 | -0.024 | -6.251 | [-0.031, -0.016] | 3.99E-09 | 1.14E-08 | AIC=2616.4; R2m=0.069; R2c=0.102 |
| EN-RAGE | -0.024 | -6.463 | [-0.031, -0.017] | 1.63E-10 | 5.62E-10 | AIC=2400.1; R2m=0.070; R2c=0.251 |
| IL-12B | -0.025 | -7.865 | [-0.031, -0.019] | 4.84E-14 | 3.03E-13 | AIC=2105.5; R2m=0.108; R2c=0.168 |
| STAMBP | -0.025 | -4.690 | [-0.036, -0.015] | 3.10E-06 | 7.14E-06 | AIC=3249.2; R2m=0.038; R2c=0.247 |
| AXIN1 | -0.029 | -4.330 | [-0.042, -0.016] | 1.62E-05 | 3.10E-05 | AIC=3695.1; R2m=0.024; R2c=0.483 |
| TWEAK | -0.030 | -15.969 | [-0.034, -0.027] | 3.13E-35 | 4.32E-34 | AIC=981.8; R2m=0.327; R2c=0.354 |
| TRANCE | -0.032 | -12.443 | [-0.037, -0.027] | 1.87E-13 | 1.07E-12 | AIC=1925.1; R2m=0.202; R2c=0.212 |
| SIRT2 | -0.033 | -4.750 | [-0.046, -0.019] | 2.31E-06 | 5.49E-06 | AIC=3810.3; R2m=0.036; R2c=0.300 |
| MCP-4 | -0.037 | -8.908 | [-0.045, -0.029] | 3.64E-18 | 3.14E-17 | AIC=2657.3; R2m=0.137; R2c=0.254 |
| CD8A | -0.051 | -12.409 | [-0.059, -0.043] | 7.29E-32 | 8.38E-31 | AIC=2650.3; R2m=0.225; R2c=0.308 |
| CXCL5 | -0.060 | -7.137 | [-0.076, -0.043] | 1.73E-12 | 7.97E-12 | AIC=4251.7; R2m=0.081; R2c=0.346 |

Supplementary Table 8. Spline models for continuous gestational age. Abbreviations: AIC, Akaike information criterion; FDR, false discovery rate; NPX, Normalized Protein eXpression.

| **Protein** | **F** | **p (uncorrected)** | **p (FDR-adjusted)** | **Predicted ΔNPX (late–early)** | **Model fit** |
| --- | --- | --- | --- | --- | --- |
| CCL28 | 566.594 | 1.76E-122 | 4.04E-121 | 3.145 | AIC=3027.6; R2m=0.334; R2c=0.334 |
| LIF-R | 697.886 | 6.03E-151 | 2.08E-149 | 2.371 | AIC=1725.4; R2m=0.512; R2c=0.526 |
| PD-L1 | 932.475 | 8.00E-202 | 5.52E-200 | 2.177 | AIC=1611.4; R2m=0.598; R2c=0.618 |
| FGF-21 | 45.152 | 8.59E-10 | 1.92E-09 | 1.238 | AIC=4751.4; R2m=0.084; R2c=0.109 |
| uPA | 535.116 | 1.17E-115 | 2.02E-114 | 1.208 | AIC=830.1; R2m=0.548; R2c=0.593 |
| LAP-TGF-β-1 | 289.270 | 2.09E-62 | 2.40E-61 | 1.083 | AIC=1617.2; R2m=0.400; R2c=0.473 |
| OPG | 237.216 | 3.81E-51 | 3.75E-50 | 1.000 | AIC=1977.4; R2m=0.236; R2c=0.239 |
| CDCP1 | 189.409 | 8.19E-41 | 7.06E-40 | 0.894 | AIC=1980.6; R2m=0.158; R2c=0.159 |
| FGF-23 | 130.941 | 3.39E-28 | 1.80E-27 | 0.789 | AIC=4277.1; R2m=0.175; R2c=0.179 |
| HGF | 102.024 | 5.70E-22 | 2.62E-21 | 0.707 | AIC=1260.4; R2m=0.150; R2c=0.189 |
| IL-17C | 32.868 | 3.43E-07 | 5.92E-07 | 0.614 | AIC=2366.0; R2m=0.070; R2c=0.138 |
| MMP-1 | 6.326 | 9.68E-02 | 1.06E-01 | 0.501 | AIC=3493.4; R2m=0.012; R2c=0.184 |
| IL-10RB | 157.903 | 5.19E-34 | 3.58E-33 | 0.474 | AIC=230.4; R2m=0.242; R2c=0.287 |
| OSM | 5.595 | 1.33E-01 | 1.43E-01 | 0.429 | AIC=3111.7; R2m=0.017; R2c=0.096 |
| IL18 | 28.402 | 2.99E-06 | 4.80E-06 | 0.381 | AIC=2055.0; R2m=0.046; R2c=0.073 |
| TGF-α | 35.268 | 1.07E-07 | 1.94E-07 | 0.352 | AIC=1124.1; R2m=0.028; R2c=0.248 |
| CCL25 | 36.116 | 7.08E-08 | 1.32E-07 | 0.315 | AIC=1939.3; R2m=0.036; R2c=0.036 |
| VEGFA | 55.849 | 4.52E-12 | 1.25E-11 | 0.311 | AIC=243.6; R2m=0.102; R2c=0.202 |
| IL-15RA | 38.711 | 2.00E-08 | 3.94E-08 | 0.307 | AIC=491.3; R2m=0.078; R2c=0.116 |
| CSF-1 | 50.551 | 6.10E-11 | 1.62E-10 | 0.237 | AIC=-392.3; R2m=0.075; R2c=0.139 |
| Flt3L | 7.236 | 6.47E-02 | 7.32E-02 | 0.221 | AIC=1193.2; R2m=0.036; R2c=0.047 |
| IL-18R1 | 21.127 | 9.91E-05 | 1.45E-04 | 0.160 | AIC=1480.5; R2m=0.047; R2c=0.071 |
| IL6 | 18.613 | 3.29E-04 | 4.54E-04 | 0.134 | AIC=2667.6; R2m=0.031; R2c=0.096 |
| IL8 | 42.559 | 3.05E-09 | 6.49E-09 | 0.116 | AIC=2404.9; R2m=0.039; R2c=0.119 |
| IL10 | 3.212 | 3.60E-01 | 3.65E-01 | 0.110 | AIC=2345.4; R2m=0.011; R2c=0.011 |
| TNF | 2.577 | 4.62E-01 | 4.62E-01 | 0.109 | AIC=1773.9; R2m=0.006; R2c=0.033 |
| CST5 | 14.450 | 2.35E-03 | 3.01E-03 | 0.030 | AIC=1809.8; R2m=0.025; R2c=0.050 |
| CCL3 | 9.086 | 2.82E-02 | 3.35E-02 | 0.020 | AIC=2471.1; R2m=0.022; R2c=0.063 |
| CXCL11 | 6.977 | 7.26E-02 | 8.08E-02 | 0.015 | AIC=3208.3; R2m=0.007; R2c=0.152 |
| CCL4 | 4.438 | 2.18E-01 | 2.24E-01 | -0.020 | AIC=2005.8; R2m=0.008; R2c=0.069 |
| FGF-19 | 5.195 | 1.58E-01 | 1.68E-01 | -0.035 | AIC=3350.5; R2m=0.014; R2c=0.017 |
| MCP-1 | 36.435 | 6.06E-08 | 1.16E-07 | -0.112 | AIC=1661.4; R2m=0.041; R2c=0.050 |
| NT-3 | 47.891 | 2.25E-10 | 5.34E-10 | -0.128 | AIC=1678.8; R2m=0.049; R2c=0.050 |
| MMP-10 | 18.810 | 2.99E-04 | 4.21E-04 | -0.156 | AIC=2090.2; R2m=0.030; R2c=0.030 |
| CXCL9 | 14.542 | 2.25E-03 | 2.93E-03 | -0.184 | AIC=2754.5; R2m=0.023; R2c=0.027 |
| ADA | 12.384 | 6.18E-03 | 7.61E-03 | -0.205 | AIC=1815.8; R2m=0.025; R2c=0.174 |
| TNFB | 11.515 | 9.24E-03 | 1.12E-02 | -0.234 | AIC=1302.7; R2m=0.013; R2c=0.027 |
| TRAIL | 45.139 | 8.65E-10 | 1.92E-09 | -0.282 | AIC=567.4; R2m=0.046; R2c=0.097 |
| TNFRSF9 | 70.946 | 2.68E-15 | 8.80E-15 | -0.294 | AIC=1067.9; R2m=0.078; R2c=0.120 |
| CCL20 | 4.749 | 1.91E-01 | 2.00E-01 | -0.297 | AIC=3247.3; R2m=0.019; R2c=0.020 |
| IL7 | 33.783 | 2.20E-07 | 3.89E-07 | -0.297 | AIC=1928.6; R2m=0.048; R2c=0.277 |
| CX3CL1 | 32.708 | 3.71E-07 | 6.25E-07 | -0.314 | AIC=1376.6; R2m=0.047; R2c=0.091 |
| DNER | 79.819 | 3.36E-17 | 1.36E-16 | -0.346 | AIC=262.4; R2m=0.117; R2c=0.138 |
| CD40 | 39.714 | 1.23E-08 | 2.49E-08 | -0.358 | AIC=1417.9; R2m=0.049; R2c=0.287 |
| SCF | 73.982 | 5.99E-16 | 2.17E-15 | -0.380 | AIC=1178.2; R2m=0.079; R2c=0.083 |
| CXCL6 | 49.398 | 1.07E-10 | 2.74E-10 | -0.387 | AIC=2904.2; R2m=0.060; R2c=0.245 |
| 4E-BP1 | 15.333 | 1.55E-03 | 2.10E-03 | -0.398 | AIC=3192.1; R2m=0.017; R2c=0.055 |
| CCL19 | 26.438 | 7.72E-06 | 1.21E-05 | -0.415 | AIC=2550.8; R2m=0.027; R2c=0.058 |
| CCL11 | 171.933 | 4.87E-37 | 3.73E-36 | -0.445 | AIC=1522.7; R2m=0.158; R2c=0.174 |
| CD6 | 31.608 | 6.33E-07 | 1.04E-06 | -0.454 | AIC=2267.9; R2m=0.019; R2c=0.532 |
| CXCL1 | 9.037 | 2.88E-02 | 3.37E-02 | -0.477 | AIC=3197.1; R2m=0.020; R2c=0.323 |
| CASP-8 | 13.898 | 3.05E-03 | 3.82E-03 | -0.507 | AIC=2990.9; R2m=0.023; R2c=0.316 |
| CD244 | 42.524 | 3.11E-09 | 6.49E-09 | -0.535 | AIC=938.1; R2m=0.069; R2c=0.142 |
| TNFSF14 | 72.423 | 1.29E-15 | 4.46E-15 | -0.538 | AIC=2061.3; R2m=0.083; R2c=0.273 |
| CD5 | 66.208 | 2.77E-14 | 8.68E-14 | -0.596 | AIC=952.0; R2m=0.069; R2c=0.299 |
| IFN-γ | 15.006 | 1.81E-03 | 2.40E-03 | -0.680 | AIC=3491.9; R2m=0.019; R2c=0.019 |
| EN-RAGE | 48.485 | 1.68E-10 | 4.14E-10 | -0.750 | AIC=2397.6; R2m=0.066; R2c=0.249 |
| MCP-2 | 61.189 | 3.27E-13 | 9.82E-13 | -0.807 | AIC=2598.9; R2m=0.072; R2c=0.118 |
| ST1A1 | 8.634 | 3.46E-02 | 3.98E-02 | -0.826 | AIC=3710.4; R2m=0.010; R2c=0.384 |
| IL-12B | 96.208 | 1.02E-20 | 4.38E-20 | -0.884 | AIC=2076.5; R2m=0.104; R2c=0.147 |
| CCL23 | 74.547 | 4.53E-16 | 1.74E-15 | -0.942 | AIC=1710.6; R2m=0.092; R2c=0.171 |
| STAMBP | 26.000 | 9.54E-06 | 1.46E-05 | -0.979 | AIC=3249.2; R2m=0.032; R2c=0.234 |
| TWEAK | 339.908 | 2.29E-73 | 3.15E-72 | -1.006 | AIC=910.7; R2m=0.326; R2c=0.367 |
| MCP-4 | 124.382 | 8.78E-27 | 4.33E-26 | -1.087 | AIC=2620.1; R2m=0.139; R2c=0.275 |
| AXIN1 | 19.541 | 2.11E-04 | 3.04E-04 | -1.127 | AIC=3698.3; R2m=0.021; R2c=0.476 |
| TRANCE | 152.507 | 7.59E-33 | 4.36E-32 | -1.174 | AIC=1917.2; R2m=0.195; R2c=0.215 |
| SIRT2 | 24.809 | 1.69E-05 | 2.54E-05 | -1.250 | AIC=3812.1; R2m=0.031; R2c=0.291 |
| CXCL5 | 57.857 | 1.69E-12 | 4.85E-12 | -1.510 | AIC=4249.3; R2m=0.094; R2c=0.367 |
| CD8A | 157.400 | 6.67E-34 | 4.19E-33 | -1.776 | AIC=2651.4; R2m=0.215; R2c=0.295 |

Supplementary Table 9. Generalized additive models for continuous gestational age. Abbreviations: AIC, Akaike information criterion; EDF, estimated degrees of freedom; FDR, false discovery rate; NPX, Normalized Protein eXpression.

| **Protein** | **EDF** | **F** | **p (uncorrected)** | **p (FDR-adjusted)** | **Predicted ΔNPX (late–early)** | **Model fit** |
| --- | --- | --- | --- | --- | --- | --- |
| CCL28 | 3.610 | 124.511 | 7.68E-97 | 1.77E-95 | 2.824 | AIC=3028.3; R2m=0.330; R2c=0.330 |
| LIF-R | 5.970 | 102.062 | 4.83E-117 | 1.67E-115 | 2.822 | AIC=1686.3; R2m=0.559; R2c=0.559 |
| PD-L1 | 5.370 | 138.443 | 1.25E-139 | 8.64E-138 | 2.407 | AIC=1598.5; R2m=0.643; R2c=0.643 |
| FGF-21 | 3.600 | 9.849 | 1.77E-08 | 3.31E-08 | 1.522 | AIC=4741.4; R2m=0.085; R2c=0.085 |
| LAP-TGF-β-1 | 4.810 | 47.897 | 1.12E-51 | 1.29E-50 | 1.305 | AIC=1600.5; R2m=0.500; R2c=0.500 |
| uPA | 4.290 | 97.216 | 2.60E-89 | 4.48E-88 | 1.251 | AIC=816.3; R2m=0.629; R2c=0.629 |
| OPG | 2.300 | 65.879 | 1.73E-37 | 1.71E-36 | 1.166 | AIC=1977.3; R2m=0.233; R2c=0.233 |
| FGF-23 | 4.650 | 20.758 | 4.39E-22 | 2.16E-21 | 1.021 | AIC=4278.0; R2m=0.172; R2c=0.172 |
| CDCP1 | 1.000 | 171.510 | 1.01E-36 | 8.69E-36 | 1.018 | AIC=1976.8; R2m=0.155; R2c=0.155 |
| IL-17C | 1.380 | 18.713 | 1.35E-06 | 2.21E-06 | 0.696 | AIC=2350.5; R2m=0.136; R2c=0.136 |
| HGF | 4.430 | 17.611 | 4.73E-18 | 2.04E-17 | 0.692 | AIC=1250.3; R2m=0.154; R2c=0.154 |
| IL-10RB | 3.300 | 36.415 | 2.70E-29 | 1.69E-28 | 0.530 | AIC=220.5; R2m=0.260; R2c=0.260 |
| IL18 | 2.260 | 8.768 | 3.57E-05 | 5.36E-05 | 0.441 | AIC=2046.2; R2m=0.072; R2c=0.072 |
| AXIN1 | 6.780 | 6.887 | 9.10E-09 | 1.79E-08 | 0.370 | AIC=3640.2; R2m=0.325; R2c=0.325 |
| VEGFA | 1.000 | 51.555 | 1.24E-12 | 3.57E-12 | 0.369 | AIC=224.7; R2m=0.202; R2c=0.202 |
| CSF-1 | 4.490 | 10.771 | 6.21E-11 | 1.53E-10 | 0.316 | AIC=-416.5; R2m=0.139; R2c=0.139 |
| IL-15RA | 2.020 | 14.421 | 4.47E-08 | 7.90E-08 | 0.311 | AIC=479.7; R2m=0.100; R2c=0.100 |
| TGF-α | 4.620 | 5.636 | 1.48E-05 | 2.27E-05 | 0.286 | AIC=1104.7; R2m=0.199; R2c=0.199 |
| IL8 | 4.460 | 9.453 | 1.77E-09 | 4.07E-09 | 0.285 | AIC=2382.0; R2m=0.082; R2c=0.082 |
| MMP-1 | 1.910 | 1.452 | 2.54E-01 | 2.74E-01 | 0.269 | AIC=3474.6; R2m=0.126; R2c=0.126 |
| CCL3 | 1.000 | 4.094 | 4.31E-02 | 4.88E-02 | 0.263 | AIC=2461.0; R2m=0.038; R2c=0.038 |
| IL-18R1 | 2.480 | 5.393 | 1.01E-03 | 1.37E-03 | 0.252 | AIC=1473.4; R2m=0.055; R2c=0.055 |
| CCL25 | 3.320 | 8.374 | 1.02E-06 | 1.75E-06 | 0.183 | AIC=1938.1; R2m=0.034; R2c=0.034 |
| Flt3L | 1.000 | 5.619 | 1.79E-02 | 2.06E-02 | 0.161 | AIC=1184.1; R2m=0.050; R2c=0.050 |
| IL7 | 5.710 | 7.639 | 6.85E-09 | 1.39E-08 | 0.151 | AIC=1890.2; R2m=0.169; R2c=0.169 |
| IL6 | 4.280 | 4.136 | 7.30E-04 | 1.01E-03 | 0.141 | AIC=2650.9; R2m=0.074; R2c=0.074 |
| STAMBP | 6.890 | 8.042 | 1.47E-10 | 3.50E-10 | 0.135 | AIC=3194.3; R2m=0.130; R2c=0.130 |
| CXCL6 | 5.190 | 10.003 | 3.31E-11 | 8.47E-11 | 0.110 | AIC=2872.5; R2m=0.122; R2c=0.122 |
| ST1A1 | 5.960 | 3.864 | 3.51E-04 | 4.94E-04 | 0.106 | AIC=3668.8; R2m=0.258; R2c=0.258 |
| OSM | 1.000 | 0.270 | 6.04E-01 | 6.13E-01 | 0.092 | AIC=3098.3; R2m=0.073; R2c=0.073 |
| CST5 | 3.030 | 3.877 | 5.38E-03 | 6.63E-03 | 0.075 | AIC=1800.2; R2m=0.024; R2c=0.024 |
| SIRT2 | 6.680 | 7.224 | 3.53E-09 | 7.86E-09 | 0.074 | AIC=3761.8; R2m=0.161; R2c=0.161 |
| IL10 | 1.600 | 0.522 | 5.49E-01 | 5.65E-01 | 0.065 | AIC=2342.9; R2m=0.010; R2c=0.010 |
| TNF | 1.440 | 0.236 | 6.87E-01 | 6.87E-01 | 0.036 | AIC=1764.6; R2m=0.025; R2c=0.025 |
| EN-RAGE | 6.400 | 10.222 | 3.27E-13 | 1.08E-12 | -0.053 | AIC=2352.1; R2m=0.230; R2c=0.230 |
| CCL4 | 1.830 | 1.321 | 2.95E-01 | 3.13E-01 | -0.085 | AIC=1992.5; R2m=0.042; R2c=0.042 |
| CD40 | 5.200 | 8.108 | 6.25E-09 | 1.31E-08 | -0.086 | AIC=1386.2; R2m=0.181; R2c=0.181 |
| CXCL11 | 2.190 | 2.064 | 1.40E-01 | 1.56E-01 | -0.087 | AIC=3190.2; R2m=0.088; R2c=0.088 |
| NT-3 | 3.670 | 10.250 | 6.11E-09 | 1.31E-08 | -0.093 | AIC=1677.4; R2m=0.047; R2c=0.047 |
| MCP-1 | 3.510 | 8.475 | 1.17E-06 | 1.98E-06 | -0.099 | AIC=1654.8; R2m=0.043; R2c=0.043 |
| CCL20 | 2.080 | 0.948 | 3.07E-01 | 3.21E-01 | -0.131 | AIC=3245.8; R2m=0.017; R2c=0.017 |
| ADA | 2.830 | 3.350 | 1.34E-02 | 1.60E-02 | -0.158 | AIC=1795.8; R2m=0.120; R2c=0.120 |
| TNFB | 2.160 | 3.651 | 1.68E-02 | 1.97E-02 | -0.175 | AIC=1296.8; R2m=0.021; R2c=0.021 |
| FGF-19 | 1.000 | 1.465 | 2.25E-01 | 2.47E-01 | -0.200 | AIC=3345.2; R2m=0.016; R2c=0.016 |
| TNFRSF9 | 3.740 | 14.717 | 3.49E-13 | 1.09E-12 | -0.226 | AIC=1058.3; R2m=0.085; R2c=0.085 |
| TRAIL | 3.190 | 10.555 | 2.34E-08 | 4.24E-08 | -0.252 | AIC=556.6; R2m=0.073; R2c=0.073 |
| CXCL9 | 2.550 | 4.105 | 5.84E-03 | 7.06E-03 | -0.273 | AIC=2751.6; R2m=0.023; R2c=0.023 |
| TNFSF14 | 5.520 | 12.436 | 8.33E-15 | 3.19E-14 | -0.280 | AIC=2032.7; R2m=0.180; R2c=0.180 |
| MMP-10 | 1.000 | 17.127 | 3.79E-05 | 5.57E-05 | -0.306 | AIC=2085.8; R2m=0.028; R2c=0.028 |
| CX3CL1 | 2.660 | 9.213 | 2.97E-06 | 4.76E-06 | -0.329 | AIC=1365.3; R2m=0.069; R2c=0.069 |
| DNER | 3.010 | 19.814 | 6.11E-15 | 2.48E-14 | -0.370 | AIC=253.6; R2m=0.159; R2c=0.159 |
| 4E-BP1 | 2.810 | 4.317 | 3.39E-03 | 4.42E-03 | -0.392 | AIC=3180.4; R2m=0.034; R2c=0.034 |
| CCL19 | 2.900 | 6.982 | 5.20E-05 | 7.47E-05 | -0.410 | AIC=2541.4; R2m=0.034; R2c=0.034 |
| SCF | 3.210 | 16.929 | 1.95E-13 | 6.74E-13 | -0.462 | AIC=1175.8; R2m=0.082; R2c=0.082 |
| CD244 | 3.500 | 10.306 | 1.00E-08 | 1.92E-08 | -0.490 | AIC=922.1; R2m=0.070; R2c=0.070 |
| CD6 | 3.370 | 7.538 | 4.55E-06 | 7.13E-06 | -0.515 | AIC=2236.9; R2m=0.455; R2c=0.455 |
| CASP-8 | 2.040 | 4.951 | 3.62E-03 | 4.62E-03 | -0.532 | AIC=2965.7; R2m=0.260; R2c=0.260 |
| CD5 | 2.750 | 18.126 | 8.84E-13 | 2.65E-12 | -0.550 | AIC=931.0; R2m=0.246; R2c=0.246 |
| CXCL1 | 1.000 | 8.638 | 3.32E-03 | 4.40E-03 | -0.552 | AIC=3169.7; R2m=0.193; R2c=0.193 |
| CCL11 | 4.660 | 29.359 | 2.17E-31 | 1.50E-30 | -0.583 | AIC=1517.7; R2m=0.167; R2c=0.167 |
| IFN-γ | 1.390 | 8.626 | 3.86E-03 | 4.85E-03 | -0.588 | AIC=3488.4; R2m=0.019; R2c=0.019 |
| MCP-2 | 3.120 | 15.243 | 7.32E-12 | 2.02E-11 | -0.774 | AIC=2586.4; R2m=0.102; R2c=0.102 |
| IL-12B | 3.480 | 22.445 | 4.67E-19 | 2.15E-18 | -0.784 | AIC=2063.4; R2m=0.100; R2c=0.100 |
| TWEAK | 6.010 | 49.524 | 1.01E-62 | 1.39E-61 | -0.813 | AIC=884.2; R2m=0.385; R2c=0.385 |
| CCL23 | 3.150 | 17.817 | 5.44E-14 | 1.98E-13 | -0.868 | AIC=1697.3; R2m=0.167; R2c=0.167 |
| MCP-4 | 3.860 | 26.349 | 2.60E-24 | 1.38E-23 | -1.001 | AIC=2599.0; R2m=0.230; R2c=0.230 |
| TRANCE | 2.570 | 45.168 | 4.66E-29 | 2.68E-28 | -1.160 | AIC=1909.1; R2m=0.210; R2c=0.210 |
| CXCL5 | 3.940 | 12.645 | 9.65E-12 | 2.56E-11 | -1.331 | AIC=4223.2; R2m=0.180; R2c=0.180 |
| CD8A | 1.720 | 71.757 | 8.75E-32 | 6.71E-31 | -1.781 | AIC=2634.2; R2m=0.223; R2c=0.223 |

Supplementary Table 10. Fetal sex interaction models. Supplementary Tables 10 and 11 show model statistics from generalized additive models (GAMs) testing for gestational age (GA) × categorical modifier (fetal sex or gravidity) interactions. Each protein was modeled with a smooth term for GA and modifier-specific smooths to test for differential associations. The F-statistic and associated numerator (Num. df) and denominator (Den. df) degrees of freedom reflect the joint test comparing the full (interaction) model to a reduced model with a single GA smooth shared across groups. Akaike information criterion values indicate overall model fit. Reported p-values correspond to the uncorrected and false discovery rate-adjusted significance levels for the joint interaction test. NA values indicate proteins for which the full and baseline models yielded indistinguishable fits, resulting in no valid comparison (i.e., no additional variance explained by the modifier-specific smooths). Abbreviations: AIC, Akaike information criterion; FDR, false discovery rate.

| **Protein** | **F** | **Num. df** | **Den. df** | **AIC** | **p (uncorrected)** | **p (FDR-adjusted)** |
| --- | --- | --- | --- | --- | --- | --- |
| CD244 | 5.130 | 4.050 | 1138.690 | 910.920 | <0.001 | 0.026 |
| LAP-TGF-β-1 | 3.410 | 4.100 | 1135.480 | 1591.330 | 0.008 | 0.266 |
| CD8A | 7.760 | 0.400 | 1141.890 | 2631.980 | 0.024 | 0.500 |
| IL-17C | 2.860 | 3.270 | 1139.570 | 2345.940 | 0.032 | 0.500 |
| IL18 | 2.570 | 3.870 | 1136.900 | 2043.140 | 0.038 | 0.500 |
| CCL19 | 3.650 | 1.060 | 1139.300 | 2539.690 | 0.054 | 0.540 |
| PD-L1 | 3.370 | 1.010 | 1138.630 | 1598.520 | 0.066 | 0.540 |
| TRANCE | 3.740 | 0.840 | 1139.600 | 1908.060 | 0.060 | 0.540 |
| NT-3 | 2.750 | 1.010 | 1141.270 | 1675.690 | 0.097 | 0.702 |
| MCP-1 | 1.880 | 3.740 | 1136.520 | 1654.100 | 0.117 | 0.758 |
| VEGFA | 0.980 | 1.000 | 1141.850 | 225.790 | 0.322 | 0.873 |
| IL6 | 1.350 | 2.470 | 1138.000 | 2655.210 | 0.260 | 0.873 |
| CXCL11 | 0.680 | 1.040 | 1140.040 | 3191.380 | 0.414 | 0.873 |
| TRAIL | 2.090 | 0.970 | 1139.280 | 556.460 | 0.149 | 0.873 |
| CXCL9 | 0.720 | 1.090 | 1140.600 | 2753.140 | 0.409 | 0.873 |
| OSM | 0.560 | 0.990 | 1141.860 | 3099.710 | 0.454 | 0.873 |
| CXCL1 | 0.380 | 0.990 | 1141.940 | 3171.320 | 0.536 | 0.873 |
| CCL4 | 0.350 | 1.100 | 1140.830 | 1994.620 | 0.574 | 0.873 |
| MCP-4 | 0.490 | 0.980 | 1138.910 | 2600.030 | 0.479 | 0.873 |
| TNFSF14 | 0.840 | 0.960 | 1138.810 | 2040.010 | 0.354 | 0.873 |
| MMP-1 | 0.400 | 0.730 | 1141.240 | 3476.000 | 0.463 | 0.873 |
| FGF-21 | 0.270 | 1.010 | 1139.180 | 4744.530 | 0.603 | 0.873 |
| IL-10RB | 0.250 | 1.000 | 1141.830 | 223.890 | 0.618 | 0.873 |
| IL-18R1 | 0.380 | 1.030 | 1139.680 | 1475.950 | 0.546 | 0.873 |
| CXCL5 | 0.250 | 1.000 | 1141.920 | 4230.100 | 0.614 | 0.873 |
| HGF | 1.000 | 1.000 | 1138.890 | 1252.220 | 0.318 | 0.873 |
| IL-12B | 0.310 | 1.020 | 1138.700 | 2064.750 | 0.580 | 0.873 |
| MMP-10 | 1.140 | 1.000 | 1147.000 | 2086.650 | 0.286 | 0.873 |
| TNF | 1.840 | 0.990 | 1141.890 | 1763.770 | 0.175 | 0.873 |
| CCL23 | 0.500 | 0.890 | 1139.240 | 1699.030 | 0.461 | 0.873 |
| CCL3 | 0.720 | 1.000 | 1141.600 | 2462.730 | 0.396 | 0.873 |
| Flt3L | 0.930 | 1.000 | 1142.250 | 1185.470 | 0.334 | 0.873 |
| CXCL6 | 1.470 | 3.530 | 1136.070 | 2876.250 | 0.215 | 0.873 |
| 4E-BP1 | 0.810 | 0.960 | 1139.530 | 3182.130 | 0.363 | 0.873 |
| CCL28 | 0.340 | 0.990 | 1144.130 | 3030.090 | 0.555 | 0.873 |
| CD40 | 0.640 | 1.000 | 1138.640 | 1389.140 | 0.424 | 0.873 |
| IFN-γ | 0.340 | 1.000 | 1143.210 | 3489.890 | 0.559 | 0.873 |
| FGF-19 | 0.700 | 1.010 | 1142.790 | 3346.970 | 0.403 | 0.873 |
| MCP-2 | 0.910 | 1.000 | 1139.360 | 2587.440 | 0.340 | 0.873 |
| CCL25 | 1.380 | 1.050 | 1144.170 | 1938.500 | 0.241 | 0.873 |
| CX3CL1 | 0.370 | 1.010 | 1139.470 | 1366.990 | 0.547 | 0.873 |
| TNFRSF9 | 0.510 | 0.990 | 1138.990 | 1059.480 | 0.474 | 0.873 |
| TWEAK | 0.740 | 1.010 | 1138.920 | 897.810 | 0.391 | 0.873 |
| CCL20 | 1.730 | 1.580 | 1146.410 | 3245.300 | 0.185 | 0.873 |
| STAMBP | 0.280 | 1.000 | 1138.640 | 3215.070 | 0.599 | 0.873 |
| TNFB | 0.610 | 1.080 | 1140.400 | 1298.980 | 0.446 | 0.873 |
| IL8 | 0.110 | 0.960 | 1138.620 | 2382.260 | 0.724 | 0.898 |
| uPA | 0.090 | 1.000 | 1138.840 | 816.810 | 0.764 | 0.898 |
| CST5 | 0.090 | 1.020 | 1139.110 | 1802.580 | 0.767 | 0.898 |
| SCF | 0.080 | 0.980 | 1140.290 | 1177.400 | 0.773 | 0.898 |
| CCL11 | 0.120 | 1.000 | 1138.440 | 1519.290 | 0.728 | 0.898 |
| LIF-R | 0.110 | 1.000 | 1138.700 | 1690.030 | 0.739 | 0.898 |
| DNER | 0.100 | 1.010 | 1139.630 | 256.110 | 0.753 | 0.898 |
| CASP-8 | 0.110 | 1.030 | 1141.030 | 2967.900 | 0.753 | 0.898 |
| ADA | 0.200 | 1.010 | 1139.630 | 1797.860 | 0.655 | 0.898 |
| CSF-1 | 0.090 | 1.000 | 1138.390 | -414.890 | 0.764 | 0.898 |
| EN-RAGE | 0.060 | 1.000 | 1138.620 | 2367.060 | 0.807 | 0.921 |
| CDCP1 | 0.010 | 0.950 | 1146.770 | 1977.680 | 0.894 | 0.965 |
| IL7 | 0.020 | 1.000 | 1138.640 | 1898.270 | 0.893 | 0.965 |
| OPG | 0.010 | 1.000 | 1142.290 | 1980.220 | 0.908 | 0.965 |
| CD6 | 0.020 | 1.010 | 1139.540 | 2239.660 | 0.876 | 0.965 |
| TGF-α | 0.010 | 1.000 | 1138.740 | 1107.600 | 0.930 | 0.965 |
| IL-15RA | 0.000 | 0.860 | 1140.560 | 481.740 | 0.965 | 0.965 |
| IL10 | 0.000 | 1.000 | 1147.000 | 2343.340 | 0.949 | 0.965 |
| ST1A1 | 0.000 | 1.000 | 1141.940 | 3683.410 | 0.960 | 0.965 |
| AXIN1 | NA | NA | NA | NA | NA | NA |
| FGF-23 | NA | NA | NA | NA | NA | NA |
| CD5 | NA | NA | NA | NA | NA | NA |
| SIRT2 | NA | NA | NA | NA | NA | NA |

Supplementary Table 11. Gravidity interaction models. Each protein was modeled with a smooth term for GA and modifier-specific smooths to test for differential associations. The F-statistic and associated numerator (Num. df) and denominator (Den. df) degrees of freedom reflect the joint test comparing the full (interaction) model to a reduced model with a single GA smooth shared across groups. Akaike information criterion values indicate overall model fit. Reported p-values correspond to the uncorrected and false discovery rate-adjusted significance levels for the joint interaction test. NA values indicate proteins for which the full and baseline models yielded indistinguishable fits, resulting in no valid comparison (i.e., no additional variance explained by the modifier-specific smooths). Abbreviations: AIC, Akaike information criterion; FDR, false discovery rate.

| **Protein** | **F** | **Num. df** | **Den. df** | **AIC** | **p (uncorrected)** | **p (FDR-adjusted)** |
| --- | --- | --- | --- | --- | --- | --- |
| CST5 | 26.500 | 0.300 | 1139.820 | 1792.200 | 0.001 | 0.050 |
| SIRT2 | 4.030 | 4.810 | 1137.890 | 3776.720 | 0.001 | 0.050 |
| ADA | 3.070 | 4.650 | 1136.000 | 1789.980 | 0.011 | 0.255 |
| CDCP1 | 3.470 | 1.720 | 1145.990 | 1973.380 | 0.038 | 0.432 |
| IL-15RA | 2.980 | 2.580 | 1138.840 | 477.030 | 0.038 | 0.432 |
| CASP-8 | 2.710 | 3.930 | 1138.140 | 2962.560 | 0.030 | 0.432 |
| LAP-TGF-β-1 | 2.360 | 1.990 | 1137.590 | 1597.440 | 0.095 | 0.467 |
| IL6 | 2.960 | 1.880 | 1138.600 | 2652.300 | 0.056 | 0.467 |
| CD6 | 2.090 | 3.410 | 1137.140 | 2236.570 | 0.092 | 0.467 |
| MMP-1 | 2.160 | 2.760 | 1139.200 | 3473.870 | 0.096 | 0.467 |
| CCL19 | 2.650 | 1.820 | 1138.540 | 2540.190 | 0.076 | 0.467 |
| CCL23 | 2.990 | 1.420 | 1138.710 | 1696.390 | 0.068 | 0.467 |
| DNER | 2.900 | 1.970 | 1138.660 | 252.440 | 0.056 | 0.467 |
| CCL20 | 2.170 | 3.550 | 1144.440 | 3243.830 | 0.078 | 0.467 |
| MMP-10 | 2.130 | 2.000 | 1146.000 | 2085.510 | 0.119 | 0.539 |
| CXCL11 | 1.810 | 2.750 | 1138.330 | 3190.110 | 0.149 | 0.545 |
| Flt3L | 1.940 | 1.660 | 1141.580 | 1184.450 | 0.152 | 0.545 |
| 4E-BP1 | 2.060 | 1.820 | 1138.680 | 3180.780 | 0.133 | 0.545 |
| FGF-19 | 1.750 | 3.620 | 1140.170 | 3345.740 | 0.143 | 0.545 |
| CD244 | 1.530 | 5.080 | 1137.670 | 925.830 | 0.178 | 0.547 |
| TNFSF14 | 1.640 | 3.400 | 1136.360 | 2039.250 | 0.171 | 0.547 |
| NT-3 | 1.640 | 2.670 | 1139.610 | 1677.040 | 0.183 | 0.547 |
| STAMBP | 1.700 | 1.900 | 1137.740 | 3213.430 | 0.185 | 0.547 |
| IL-18R1 | 1.620 | 1.950 | 1138.760 | 1474.860 | 0.199 | 0.565 |
| IL8 | 1.540 | 2.140 | 1137.430 | 2380.910 | 0.214 | 0.576 |
| TNFB | 1.520 | 1.940 | 1139.540 | 1298.300 | 0.220 | 0.576 |
| LIF-R | 1.280 | 2.040 | 1137.670 | 1689.500 | 0.278 | 0.675 |
| IL-10RB | 1.270 | 5.020 | 1137.810 | 224.560 | 0.274 | 0.675 |
| CD8A | 1.200 | 1.400 | 1140.890 | 2635.420 | 0.290 | 0.680 |
| SCF | 1.140 | 2.690 | 1138.570 | 1177.230 | 0.328 | 0.697 |
| IL18 | 1.120 | 1.960 | 1138.810 | 2048.040 | 0.327 | 0.697 |
| CD5 | 1.190 | 2.810 | 1137.690 | 932.360 | 0.311 | 0.697 |
| TGF-α | 0.970 | 2.000 | 1137.740 | 1107.630 | 0.381 | 0.728 |
| TNF | 0.940 | 1.860 | 1141.020 | 1765.430 | 0.386 | 0.728 |
| IFN-γ | 1.040 | 2.200 | 1142.020 | 3490.270 | 0.358 | 0.728 |
| CX3CL1 | 0.870 | 1.400 | 1139.070 | 1366.860 | 0.385 | 0.728 |
| IL7 | 0.860 | 2.010 | 1137.630 | 1898.510 | 0.425 | 0.729 |
| CXCL9 | 0.840 | 2.300 | 1139.390 | 2754.370 | 0.445 | 0.729 |
| CCL11 | 0.920 | 2.800 | 1136.640 | 1519.850 | 0.425 | 0.729 |
| CCL28 | 0.800 | 1.990 | 1143.130 | 3030.820 | 0.447 | 0.729 |
| TWEAK | 1.010 | 3.750 | 1136.180 | 899.410 | 0.398 | 0.729 |
| ST1A1 | 0.800 | 2.000 | 1140.940 | 3683.790 | 0.450 | 0.729 |
| VEGFA | 0.780 | 2.000 | 1140.840 | 227.150 | 0.461 | 0.729 |
| CCL3 | 0.730 | 2.080 | 1140.510 | 2464.000 | 0.490 | 0.757 |
| CXCL1 | 0.660 | 2.000 | 1140.930 | 3172.350 | 0.515 | 0.762 |
| TNFRSF9 | 0.670 | 1.990 | 1137.990 | 1060.510 | 0.511 | 0.762 |
| OPG | 0.680 | 2.560 | 1140.730 | 1981.410 | 0.544 | 0.776 |
| FGF-23 | 0.390 | 1.070 | 1142.760 | 4277.990 | 0.548 | 0.776 |
| IL-17C | 0.560 | 1.990 | 1140.850 | 2352.590 | 0.568 | 0.783 |
| PD-L1 | 0.550 | 1.960 | 1137.670 | 1602.720 | 0.576 | 0.783 |
| uPA | 0.450 | 1.960 | 1137.880 | 817.880 | 0.634 | 0.798 |
| MCP-1 | 0.410 | 1.570 | 1138.690 | 1657.000 | 0.613 | 0.798 |
| HGF | 0.460 | 1.890 | 1137.990 | 1254.080 | 0.624 | 0.798 |
| CCL25 | 0.460 | 1.870 | 1143.350 | 1940.740 | 0.621 | 0.798 |
| CCL4 | 0.330 | 2.220 | 1139.710 | 1996.380 | 0.738 | 0.912 |
| TRAIL | 0.220 | 1.900 | 1138.350 | 559.920 | 0.791 | 0.945 |
| FGF-21 | 0.180 | 1.670 | 1138.520 | 4745.510 | 0.792 | 0.945 |
| TRANCE | 0.180 | 1.760 | 1138.680 | 1912.580 | 0.809 | 0.948 |
| CXCL6 | 0.170 | 2.030 | 1137.570 | 2879.120 | 0.843 | 0.955 |
| CD40 | 0.190 | 2.020 | 1137.620 | 1391.360 | 0.831 | 0.955 |
| OSM | 0.090 | 1.960 | 1140.880 | 3102.010 | 0.914 | 0.972 |
| EN-RAGE | 0.090 | 2.010 | 1137.610 | 2368.920 | 0.915 | 0.972 |
| MCP-2 | 0.080 | 1.860 | 1138.500 | 2589.860 | 0.908 | 0.972 |
| CSF-1 | 0.100 | 2.030 | 1137.360 | -413.080 | 0.909 | 0.972 |
| IL10 | 0.060 | 2.000 | 1146.000 | 2345.220 | 0.944 | 0.988 |
| MCP-4 | 0.020 | 2.020 | 1137.870 | 2602.510 | 0.977 | 0.991 |
| IL-12B | 0.020 | 1.840 | 1137.880 | 2066.600 | 0.969 | 0.991 |
| CXCL5 | 0.010 | 2.000 | 1140.920 | 4232.330 | 0.992 | 0.992 |
| AXIN1 | NA | NA | NA | NA | NA | NA |

Supplementary Table 12. Maternal age interaction models. This table shows model statistics from generalized additive models testing for continuous modification of gestational age (GA) trajectories by maternal age at blood draw. Each protein was modeled with main-effect smooths for GA and maternal age and a tensor-product interaction smooth to capture continuous modification of GA–protein relationships by maternal age. The estimated degrees of freedom and F-statistic describe the shape and strength of the interaction smooth, respectively, and Akaike information criterion values indicate overall model fit. Reported p-values correspond to the uncorrected and false discovery rate-adjusted significance levels for the maternal age interaction term. Abbreviations: AIC, Akaike information criterion; EDF, estimated degrees of freedom; FDR, false discovery rate.

| Protein | EDF | F | AIC | p (uncorrected) | p (FDR-adjusted) |
| --- | --- | --- | --- | --- | --- |
| TGF-α | 1.000 | 8.150 | 1098.960 | 0.004 | 0.262 |
| IL10 | 7.690 | 2.190 | 2331.560 | 0.011 | 0.262 |
| TNF | 1.000 | 6.420 | 1760.690 | 0.011 | 0.262 |
| VEGFA | 2.180 | 2.860 | 221.070 | 0.036 | 0.452 |
| uPA | 1.000 | 3.990 | 813.060 | 0.046 | 0.452 |
| TNFSF14 | 3.940 | 2.270 | 2032.760 | 0.035 | 0.452 |
| HGF | 1.000 | 4.200 | 1248.370 | 0.041 | 0.452 |
| TRAIL | 1.000 | 3.480 | 555.870 | 0.062 | 0.473 |
| OSM | 1.600 | 2.550 | 3096.500 | 0.094 | 0.473 |
| CD6 | 1.480 | 2.140 | 2235.670 | 0.080 | 0.473 |
| IL-10RB | 1.630 | 2.590 | 218.750 | 0.092 | 0.473 |
| Flt3L | 2.370 | 2.080 | 1182.360 | 0.093 | 0.473 |
| DNER | 4.650 | 1.840 | 252.220 | 0.089 | 0.473 |
| CSF-1 | 3.450 | 1.830 | -419.140 | 0.096 | 0.473 |
| EN-RAGE | 1.830 | 2.050 | 2363.110 | 0.103 | 0.474 |
| IL8 | 1.000 | 1.820 | 2381.120 | 0.178 | 0.543 |
| CDCP1 | 1.000 | 1.470 | 1977.180 | 0.225 | 0.543 |
| AXIN1 | 3.440 | 1.730 | 3652.490 | 0.127 | 0.543 |
| CST5 | 1.000 | 1.430 | 1801.940 | 0.231 | 0.543 |
| CXCL1 | 5.270 | 1.320 | 3168.590 | 0.233 | 0.543 |
| FGF-23 | 1.000 | 2.030 | 4276.180 | 0.154 | 0.543 |
| FGF-21 | 1.470 | 2.470 | 4743.640 | 0.147 | 0.543 |
| IL-18R1 | 4.340 | 1.700 | 1472.010 | 0.195 | 0.543 |
| CXCL6 | 1.650 | 1.470 | 2875.280 | 0.217 | 0.543 |
| SIRT2 | 4.790 | 1.530 | 3775.140 | 0.152 | 0.543 |
| IFN-γ | 1.000 | 1.840 | 3488.130 | 0.176 | 0.543 |
| MCP-2 | 1.000 | 1.410 | 2586.770 | 0.236 | 0.543 |
| STAMBP | 5.010 | 1.480 | 3211.740 | 0.169 | 0.543 |
| ADA | 2.860 | 1.480 | 1794.440 | 0.202 | 0.543 |
| TNFB | 1.510 | 2.050 | 1296.640 | 0.205 | 0.543 |
| CD5 | 1.210 | 0.960 | 931.080 | 0.267 | 0.576 |
| TNFRSF9 | 1.000 | 1.230 | 1057.610 | 0.267 | 0.576 |
| IL7 | 5.240 | 1.080 | 1898.090 | 0.378 | 0.602 |
| IL6 | 1.330 | 1.430 | 2656.200 | 0.333 | 0.602 |
| CXCL11 | 1.000 | 0.910 | 3191.120 | 0.340 | 0.602 |
| CXCL9 | 1.320 | 0.650 | 2751.500 | 0.366 | 0.602 |
| MCP-4 | 1.010 | 0.760 | 2599.800 | 0.384 | 0.602 |
| TRANCE | 1.810 | 0.970 | 1911.170 | 0.357 | 0.602 |
| CCL3 | 1.000 | 0.800 | 2463.660 | 0.369 | 0.602 |
| 4E-BP1 | 3.820 | 1.090 | 3182.040 | 0.361 | 0.602 |
| FGF-19 | 1.000 | 0.850 | 3346.590 | 0.357 | 0.602 |
| TWEAK | 4.560 | 1.060 | 899.020 | 0.381 | 0.602 |
| CCL20 | 4.570 | 1.100 | 3245.420 | 0.342 | 0.602 |
| ST1A1 | 3.710 | 1.060 | 3681.940 | 0.382 | 0.602 |
| LIF-R | 1.000 | 0.680 | 1689.120 | 0.410 | 0.629 |
| LAP-TGF-β-1 | 2.550 | 0.750 | 1599.380 | 0.444 | 0.645 |
| CCL4 | 1.000 | 0.610 | 1994.000 | 0.435 | 0.645 |
| IL18 | 1.000 | 0.570 | 2047.270 | 0.451 | 0.645 |
| IL-12B | 1.000 | 0.480 | 2064.050 | 0.486 | 0.645 |
| MMP-10 | 2.630 | 0.650 | 2086.790 | 0.470 | 0.645 |
| CASP-8 | 4.320 | 0.900 | 2968.700 | 0.485 | 0.645 |
| CX3CL1 | 1.000 | 0.520 | 1368.100 | 0.472 | 0.645 |
| CD40 | 1.600 | 0.790 | 1389.770 | 0.499 | 0.650 |
| MMP-1 | 1.740 | 0.530 | 3476.390 | 0.541 | 0.692 |
| CD244 | 1.000 | 0.240 | 923.750 | 0.627 | 0.713 |
| IL-17C | 2.220 | 0.620 | 2353.040 | 0.613 | 0.713 |
| SCF | 1.340 | 0.620 | 1177.190 | 0.611 | 0.713 |
| CCL11 | 1.990 | 0.840 | 1518.900 | 0.620 | 0.713 |
| IL-15RA | 1.000 | 0.290 | 481.240 | 0.591 | 0.713 |
| PD-L1 | 1.000 | 0.290 | 1601.540 | 0.591 | 0.713 |
| CCL23 | 2.670 | 0.680 | 1700.090 | 0.630 | 0.713 |
| MCP-1 | 1.970 | 0.360 | 1658.080 | 0.689 | 0.739 |
| CCL19 | 2.090 | 0.430 | 2545.610 | 0.677 | 0.739 |
| CCL25 | 1.460 | 0.400 | 1940.100 | 0.697 | 0.739 |
| NT-3 | 1.000 | 0.180 | 1675.710 | 0.676 | 0.739 |
| CXCL5 | 3.420 | 0.640 | 4226.420 | 0.721 | 0.754 |
| CD8A | 1.400 | 0.370 | 2636.730 | 0.736 | 0.758 |
| CCL28 | 1.580 | 0.170 | 3030.720 | 0.838 | 0.850 |
| OPG | 1.000 | 0.010 | 1977.530 | 0.920 | 0.920 |
